# The Development of Synapses in Mouse and Macaque Primary Sensory Cortices

**DOI:** 10.1101/2023.02.15.528564

**Authors:** Gregg Wildenberg, Hanyu Li, Narayanan Kasthuri

## Abstract

We report that the rate of synapse development in primary sensory cortices of mice and macaques is unrelated to lifespan, as was previously thought. We analyzed 28,084 synapses over multiple developmental time points in both species and find, instead, that net excitatory synapse development of mouse and macaque neurons primarily increased at similar rates in the first few postnatal months, and then decreased over a span of 1-1.5 years of age. The development of inhibitory synapses differed qualitatively across species. In macaques, net inhibitory synapses first increase and then decrease on excitatory soma at similar ages as excitatory synapses. In mice, however, such synapses are added throughout life. These findings contradict the long-held belief that the cycle of synapse formation and pruning occurs earlier in shorter-lived animals. Instead, our results suggest more nuanced rules, with the development of different types of synapses following different timing rules or different trajectories across species.

## 1 Introduction

A central feature of the development of neurons in mammalian cortex is the reapportionment of synapses over post-natal life with little change in numbers of neurons [1, 2, 3, 4, 5]. Evidence for synaptic re-arrangements have been found in the cortices of nearly all mammals studied [6, 7, 8, 9, 10, 11, 12, 13, 14, 15, 16] and the mechanisms of these re-arrangements are considered by some to underlie learning and memory in the adult brain [17, 18, 19]. Failures of these mechanisms have been implicated in variety of mental disorders including Autism Spectrum Disorders (ASD) and Schizophrenia [20, 21, 22, 23, 24].

However, despite its prevalence and importance, the mechanisms of synaptic development in mammalian cortex remain obscured. One option is to leverage the aforementioned remarkable homology of cortex across species, detailing cortical synaptic development across phylogeny in order to identify which mechanism are conserved and which have been altered by evolution and to what potential effect. But, until recently, the unambiguous detailing of synapses over large volumes was hard. As a result most studies of synaptic development, particualry across species and development, use a variety of approaches: optical reconstructions of axonal arbors [25, 26], paired recordings [27], trans-synaptic labeling with radioactive amino acids or modified Rabies viruses [28, 29, 30], immunohistochemistry [31], expression of genes associated with synaptic development [32, 33], and bulk synapse density measurements using single-section electron microscopy [8, 34, 35, 36, 37] to name a few. It can be difficult to quantitatively relate the results from one approach to another. Second, most approaches are indirect measures of connectivity with unspecified false positive and negative rates. For example, trans-synaptic approaches only label a small fraction of connections on individual neurons, and it is not clear whether this sparse labeling is unbiased. Synapse density measurements using single-section electron microscopy do not account for changes in neuronal geometry, such as the growth of dendrites, or variability along different parts of neurons or in different classes of synapses.

Thus we asked, what could be revealed with a more complete and unbiased reckoning of synaptic development, using the same approach to identify neuronal connections in two well studied species, *Mus musculus* (mouse) and *Macaca mulatta* (rhesus macaque or primate). We used large volume serial EM (”connectomics”) to reconstruct excitatory and inhibitory connections onto excitatory neurons from multiple cortical regions (S1 and V1 in mice and V1 in primates), from multiple cortical layers (Layers 2/3 and 4), across multiple time points post birth (p7 to p523 in mice, p7 to p3000 in primate), and across multiple animals (n= 11 mice and n=3 primates), using a combination of publicly available and newly collected data sets. We chose EM as it remains the ’gold standard’ for detailing neuronal connections and recent connectomic reports have revealed species differences [38, 39] and developmental differences in connectivity [40]. Finally, we developed an algorithmic pipeline on national lab supercomputers for automated segmentation and ‘saturated’ tracing of neurons to verify these results [41, 42].

We expected that mouse neurons would complete synaptic re-arrangements faster than primate neurons for several reasons:

- Mice live for *≈*2 years while rhesus macaques live for *≈*25 years [43, 44].
- Functional milestones correlated to synaptic re-arrangements, often termed ’critical periods’, occur earlier in mice than in macaques. For example, the period of ocular dominance formation seems to ’close’ earlier in animals with shorter lifespans, such as mice [45], compared to animals with longer lifespans (e.g., primates) [46, 47].
- A fundamental principle of modern biology is that changes in the timing and rate of developmental programs drive evolutionary adaptations [48, 49, 50], termed heterochrony.

However, we found instead:

- Little evidence for the net pruning of excitatory connections on excitatory neurons across S1 and V1 cortex in the first months of early mouse postnatal life before, during, or after the critical period. Instead, net excitatory synapse numbers grow until adulthood and then decline in older mice. Primate neurons develop excitatory synapses at nearly the same rate in absolute time as mice (e.g., synaptic development does not seem normalized with lifespan.)
- The cycle of inhibitory synapse development qualitatively differed across species. Net inhibitory synapse number increased on primate neurons for the first 75 days of life and then pruned to ≈40% of peak numbers at age p3000, with an increase in synaptic area of the remaining inputs. Net somatic innervation of mouse excitatory soma, however, continued to increase throughout the animal’s life, revealing species specific differences in inhibitory, but not excitatory, synapse development.

## 2 Results

### 3.1 Net excitatory synapse formation in the developing V1 and S1 of mice and V1 of primate

We first analyzed synaptic density on excitatory neurons in L2/3 of the primary visual cortex (V1) of mice and primates. We defined synapses as locations where an identified axon was physically proximate to an identified dendrite or soma, containing a dark contrasted postsynaptic density (PSD) and pre-synaptic axonal bouton with a vesicle cloud containing greater than ≈30-40 vesicles. We classified synapses as excitatory or inhibitory on the basis of whether they synapse onto spines versus shafts and somata, respectively [51, 38, 52, 53]. We started our first reconstructions at ≈p6 in the mouse, a time when in other systems (e.g., the developing neuromuscular junction (NMJ) and autonomic ganglia of the peripheral nervous system (PNS)), individual post-synaptic neurons receive numerous pre-synaptic inputs [1, 2, 3]. Moreover, the first post-natal week in mice is prior to eye opening (≈p14) [54, 55], a period that has been reported to contain abundant, albeit potentially weak, synaptic connections amongst cortical neurons [56, 57, 58].

Our first surprise was a near absence of synapses in the early postnatal life of the mouse. At p6, mouse excitatory neurons in L2/3 and L4 were nearly absent of synaptic innervation (e.g., across two reconstructed neurons totalling ≈125 µm of dendrite and complete soma, we found *zero* excitatory spine synapses, 89 dendritic filopodia, 1 somatic synapse, and 8 dendritic shaft synapses) **(Fig.1a, top panel, and bottom inset)**. We next evaluated spine and shaft synapse density across randomly sampled dendrites of varying diameter and orientation (e.g., dendrites likely from different parts of a neuron’s dendritic tree) to account for variability across neurons and to ensure our analyses are not biased, for example, towards proximal dendrites. Consistent with our initial observations, we found mouse excitatory neurons at this age were sparsely innervated regardless of dendrite diameter or orientation (mean±sem excitatory synapse density/µm of dendrite, mouse p6 L2/3 = 0.01±0.007, L4 = 0.02±0.008, n=20, 10µm dendrite fragments/dataset) (**Fig.1b, red lines, zoomed right inset**). In fact, of the 40 randomly sampled 10µm dendrite fragments in p6 mouse excitatory neurons in L2/3 and L4, only 6, contained spine synapses and in most cases only one.

**Figure 1.**
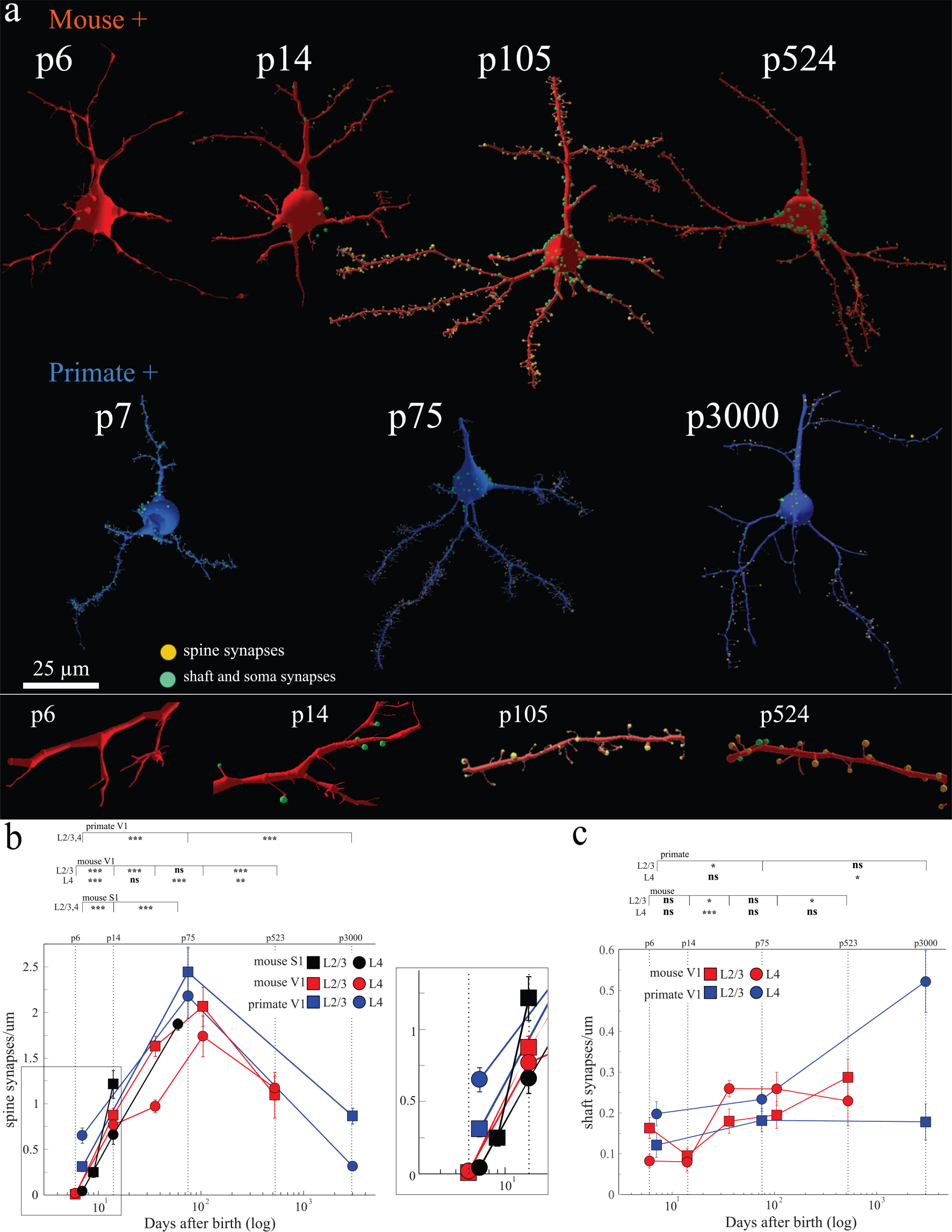
Isochronic development of excitatory synapses in primate and mouse cortex. a, Shown are representative reconstructions of V1 mouse (top, red) and primate (bottom, blue) excitatory (+) neurons at the noted postnatal (p) days. Excitatory spine synapses = orange dots, inhibitory shaft and somatic synapses = green dots. *below:*zoom-ins of mouse excitatory dendrites from ages p7, p14, p104, and p524, left to right. **b,** Scatter plot of x:postnatal days after birth (log) and y:average spine synapse density in synapses/µm. Top line postnatal (p) days. Squares = L2/3, circles = L4, black = mouse S1, red = mouse V1, blue = primate V1. *Right*: close-up of earliest data points: p6-p14 mouse and p7 primate. **c,** Scatter plot of x:postnatal days after birth (log) and y:average dendrite shaft synapses/µm. Top line postnatal (p) days. Two-sample Mann-Whitney U test, ns = P >0.05, * = P < 0.05, ** = P < 0.01, *** = P < 0.001 shown for pairwise comparisons between adjacent ages in each plot. See Supplementary Table 1 for numerical summary and supplementary files for all pairwise p-values. Lines that connect datapoints in scatter plots (b,c) are for visualization purposes only and do not represent fitted curves. Scale bar = 25 µm (a). Mouse S1 analysis from publicly available datsets: p9 and p14 [40] and p60 [118]. **(b),** mean±sem spine synapses/µm; mouse V1 L2/3 | L4, p6 = 0.007 ± 0.0065 | 0.02 ± 0.008, p14 = 0.87 ± 0.08 | 0.77 ± 0.09, p36 = 1.63 ± 0.11 | 0.97 ± 0.07, p105 = 2.1 ± 0.21 | 1.7 ± 0.22, p523 = 1.1 ± 0.3 | 1.2 ± 0.13; mouse S1 L2/3 p9 = 0.25 ± 0.04, p14 = 1.2 ± 0.15, L4 p7 = 0.04 ± 0.01, p14 = 0.66 ± 0.1, p60 = 1.87 ± 0.06; primate V1 L2/3 | L4 p7 = 0.31 ± 0.05 | 0.65 ± 0.08, p75 = 2.44 ± 0.27 | 2.2 ± 0.24, p3000 = 0.86 ± 0.09 | 0.32 ± 0.03, n = 20, 10 µm dendrite fragments/dataset and 1 animal/dataset. **(c),** mean±sem shaft synapses/µm; mouse V1 L2/3 | L4, p6 = 0.16 ± 0.03 | 0.08 ± 0.02, p14 = 0.1 ± 0.02 | 0.08 ± 0.02, p36 = 0.18 ± 0.03 | 0.26 ± 0.02, p105 = 0.20 ± 0.03 | 0.26 ± 0.04, p523 = 0.29 ± 0.04 | 0.23 ± 0.06; primate V1 L2/3 | L4, p7 = 0.12 ± 0.03 | 0.20 ± 0.03, p75 = 0.18 ± 0.03 | 0.23 ± 0.03, p3000 = 0.18 ± 0.04 | 0.52 ± 0.08, n = 20, 10 µm dendrite fragments/dataset and 1 animal/dataset.

Primate neurons at p7 also showed sparse synaptic innervation relative to adults [38]. The primate p7 neuron reconstructed in **Fig.1a** had 215 total synapses (i.e., spine, shaft, and soma) and 92 filopodia over ≈100 µm of dendritic tree reconstructed. Interestingly, while still lower than adult primate neurons, p7 primate excitatory neurons were qualitatively more spinous than similarly aged mouse neurons, with a mixture of filopodia and fully formed spine synapses (92 filopodia, 82 spine synapses). Excitatory neurons in p7 primates had 40x more spine synapses than p6 mice (e.g., mouse p6 vs primate p7; L2/3 p=2.86e-7, L4 p=3.8e-7). The few spine synapses that could be identified in mouse p6 neurons appeared fully formed, (i.e., a clear PSD and numerous synaptic vesicles, similar to primate p7 neurons) **(Supplementary Fig.1a)**. Finally, we found a slight correlation between dendrite diameter and synaptic density but the effect size was small and not significantly different **(Supplementary Fig.2)**.

**Figure 2.**
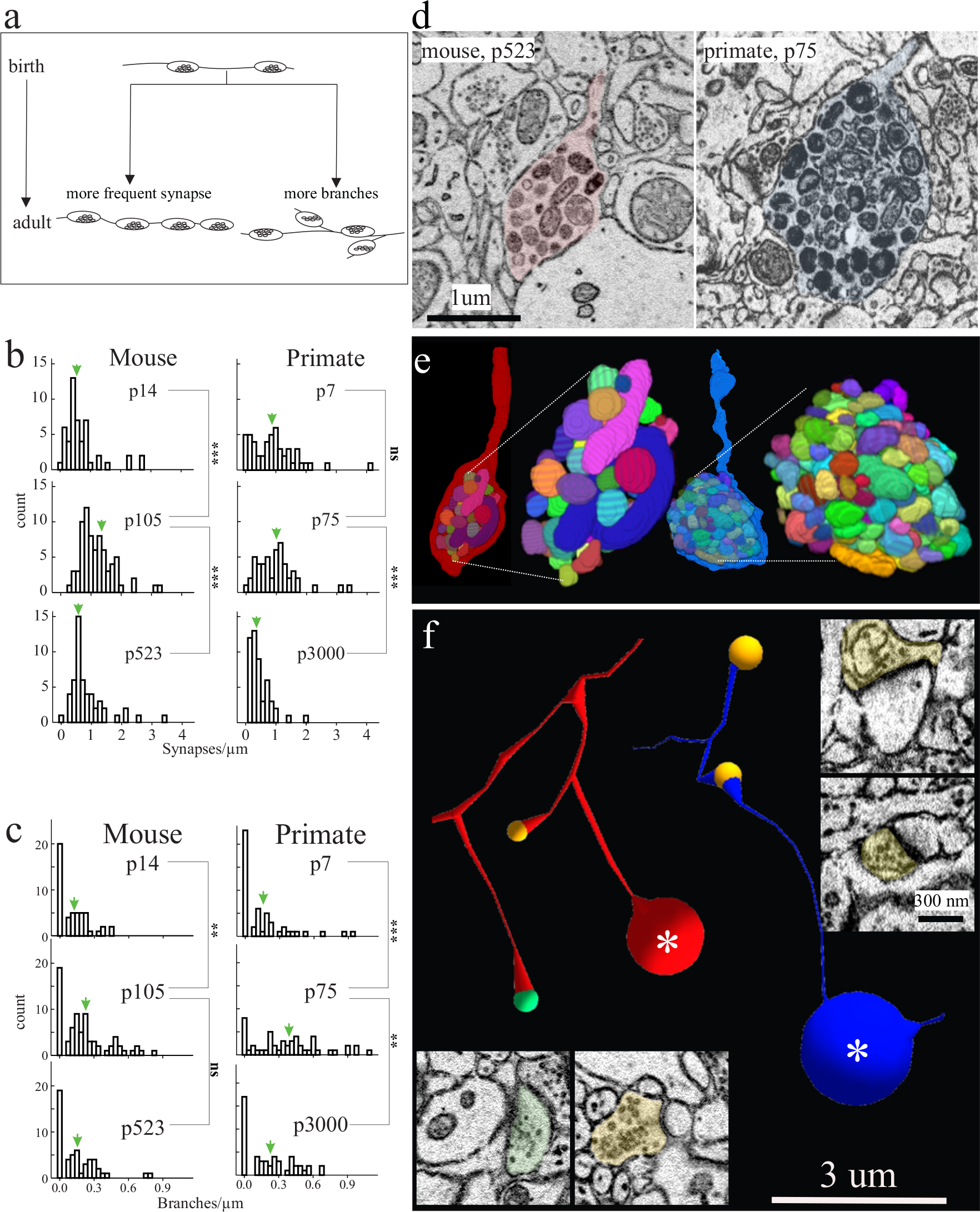
Excitatory axon development in mouse and primate. **a**, cartoon depicting hypothetical models of excitatory axon development: *left*, axons increase their synapse frequency and/or *right*, axons make more branches. **b,c** histograms of the number of synapses/µm and branches/µm, respectively, of excitatory axons at different ages in mouse (left) and primate (right) in V1 L2/3. Green arrows indicate the ≈mean. **d,** Single 2D EM images and **e**, 3D reconstructions of a representative terminal axon retraction bulbs in mouse V1, p523 (left) and primate V1, p75 (right). 3D reconstructions show individually coloured mitochondria contained within the retraction bulb. **f,** Skeleton reconstructions of mouse (red) and primate (blue) axons containing terminal retraction bulbs (asterisk) (from **d**) and spine (orange circle) or shaft (green circle) synapses. *Insets*: 2D EM images of spine and shaft synapses made by the primate (right) and mouse (bottom) axon containing a retraction bulb. Two-sample Mann-Whitney U test, ns = P > 0.05, * = P < 0.05, ** = P < 0.01, *** = P < 0.001 shown for pairwise comparisons between adjacent ages in each plot. See Supplementary Table 2 for numerical summary and supplementary files for all pairwise p-values. Scale bar = 1 µm (d), 3 µm (f) and 300 nm (f insets). **b,** mean±sem synapses/µm; mouse p14 = 0.71 ± 0.09, p105 = 1.2 ± 0.07, p523 = 0.86 ± 0.09, primate p7 = 0.93 ± 0.1, p75 = 1.0 ± 0.09, p3000 = 0.46 ± 0.05. **c,** mean±sem branches/µm; mouse p14 = 0.12 ± 0.02, p105 = 0.22 ± 0.02, p523 = 0.15 ± 0.02, primate p7 = 0.16 ± 0.03, p75 = 0.38 ± 0.04, p3000 = 0.22 ± 0.03, n = 50-60 axons/dataset.

The differences in early innervation of macaque and mouse cortical neurons prompted us to ask how synaptic densities changed over development. For example, the higher densities in primates p7 neurons could suggest that these neurons start synaptic pruning earlier while the lower densities in p6 mouse could suggest a period of synapse formation followed by pruning. Thus, we extended our analyses across multiple time points in L2/3 and L4 across both species, including time points covering the ’critical period’ in mouse cortex (i.e., p14 to p36) [45]. We found instead a seemingly monotonic increase in synapse density in both macaques and mice over the first months of postnatal life **(Fig.1b)**. As expected, primate neurons peaked in spine synapse density at age p75 [8] relative to both p7 and p3000 ages. Surprisingly, mouse excitatory neurons showed only a modest increase in the number of spine synapses from p6 to p14 (mean±sem spine synapses/µm V1 L2/3 | L4: p6 = 0.007±0.0065 | 0.02±0.008, p14 = 0.87±0.08 | 0.77±0.09, p = 1.1e-8 | 5e-8) and such densities continued to increase to age p105, peaking around the same absolute time as the primate. Like mouse V1, we next found mouse L2/3 and L4 neurons in the primary somatosensory cortex (S1) showed a similar sparsity at early postnatal life and a steady, seemingly monotonic, increase in the total number of excitatory synapses across postnatal life **(Fig.1b, black lines)**, suggesting that, net synaptic growth over postnatal development was not unique to mouse V1. Overall, spine synapse density was broadly lower in baby and juvenile mice (e.g., mean±sem spine synapses/µm for V1, L2/3: p6=0.01±0.007, p14 =0.87±0.08, p36=1.63±0.11 versus p105=2.1±0.21), whereas juvenile primates had a higher spine synapse density compared to the adult (e.g., mean±sem spine synapses/µm for V1, L2/3: p75 = 2.44±0.27 vs. p3000 = 0.86±0.09). Finally, the similarity in the rise of excitatory synaptic density across mice and primates in absolute days prompted us to ask whether mouse neurons would show evidence of net pruning, later life, during a time frame, in absolute days, when primates also showed synaptic pruning. Thus, we analyzed similar synaptic density measurements in older mice, p523 or ≈1.5 years of age. Indeed, we found that excitatory synaptic density drops in older mice **(Fig.1b)**, as previously reported in other brain regions and layers [59]. We found that, with synaptic density estimates from older mice added to the ’synaptic development’ curve, both primate and mouse neurons added and pruned excitatory synapses at similar times in absolute days one ’clock’ for synaptic development across species with disparate lifespans. In contrast, we observed a subtle, though at times not statistically significant, increase in the number of inhibitory shaft inputs onto the same randomly sampled dendrite shafts across all V1 datasets (**Fig.1c**, and **Supplementary Fig.1b**) consistent with previous reports on postnatal changes of inhibitory shaft synapses in mouse S1 [40]. We conclude:

- Little evidence of supernumerary increases in excitatory spine synapses followed by pruning during neonatal life in mouse.
- Rather, the rise and fall of excitatory spine synapses occurs at approximately the same absolute time after birth in both species.
- In both species, the frequency of inhibitory shaft synapses appears to slowly rise over the life of the animal.

### 3.2 Excitatory axon development

We next asked how excitatory axons changed during the periods of increasing and decreasing spine synaptic density we observed. We focused these analyses on L2/3 since developmental changes in spine synapse densities were equivalent between L2/3 and L4 in both species (i.e., Figure 1). We considered two, but not mutually exclusive, mechanisms: 1) existing axons could increase their synapse frequency and/or 2) axons could locally branch more (**Fig.2a**). We traced identified excitatory axons and annotated every axonal synapse and branch point made in our volumes at ages in mice and primates with large increases in net synapses (p14 to p105 in mouse and p7 to p75 in primates). Synapse density along axons increased over this period in both species **(Fig.2b)** (mean±sem synapses/µm for mouse: p14=0.71±0.09, p105=1.2±0.07, p=8.5e-9 and primate: p7=0.93±0.1, p75=1.0±0.09, p=0.38, n=50 axons/dataset). However the increase in synaptic density on axons could not account completely for the 2.4x and 7.9x increased number of synapses for mouse and primate, respectively. Thus, we asked whether local axonal branching could also contribute to increased synapse numbers. Indeed, we found that axons branched more over this period and in both species and seemed to be due to a few axons increasing branch numbers dramatically **(Fig.2c)** (mean±sem branches/µm for mouse: p14=0.12±0.02, p105=0.22±0.02, p=7.3E-3 and primate: p7=0.16±0.03, p75 = 0.38±0.04, p=1.9e-5, n=50 axons/dataset). Thus, both mechanisms of increased synapse density and localized branching of excitatory axons appear to coincide with increases in excitatory spine synapse density along neurons (i.e., Figure 1).

In both species, we find multiple examples of axons ending in large bulb-like structure reminiscent of axon retraction bulbs during periods of synaptic pruning in the mouse developing neuromuscular junction (NMJ) [60, 61]: a sudden swelling at the end of an axon containing dense tortuously packed mitochondria (**Fig.2d-e**). Furthermore, we found numerous examples where a branch of an axon would end in such a bulb while other branches of the same axon made clear synapses on postsynaptic targets **(Fig.2f)**. Bulbs were most notable in ages mouse p523 and primate p75, the ages experiencing the most relative synaptic change, however, we were also able to identify the rare bulb in mouse p105 and primate p3000 (not shown). In both species, periods of synaptic pruning were associated with a reduction in synapse density along axons and decreased branching **(Fig.2b-c)** (i.e., synaptic pruning likely occurs for both *en passant* and *terminal* synapses) (mean±sem synapses/µm for mouse: p105 = 1.2±0.07, p523 = 0.86±0.09, p=1.6e-5 and primate p75 = 1.0±0.09, p3000 = 0.46±0.05, p=2.7e-7, mean±sem branches/µm for mouse: p105 = 0.22±0.02, p523 = 0.15±0.02, p=7.0e-2 and primate p75 = 0.38±0.04, p3000 = 0.22±0.03, p=4.7e-3, n=50 axons/dataset).

### 3.3 The Development of Somatic Inhibition Across Species

We next examined the development of somatic innervation of excitatory neurons [62] in these two species. We restricted our analyses to somatic innervation of excitatory neurons which multiple reports in both mice and primates have determined are almost exclusively made by PV interneurons [63, 62, 64]. As we captured complete soma in our EM datasets, we assessed the total somatic synaptic contribution by PV innervation. First, similar to excitatory spine synapses, we find a complete sparsity of somatic innervation shortly after birth **(Fig.3a, and see Supplementary Fig.1c)**. For example, the p7 primate excitatory soma in Figure 3a had 24 synapses, and remarkably, the p7 mouse soma only had one, far less than what we and others have reported on adult somata from the same layers and regions [38, 65, 66]. Over development, we find primarily addition of so-matic synapses from p6 to p105 in mice and from p7 to p75 in macaques **(Fig.3b)** (mean±sem # synapses/soma; mouse L2/3 | L4, p6 = 1.2±0.3 | 2.2±1.0, p14=25.8±3.5 | 11.3±1.4, p36=66±3.8 | 38±0.82, p105=72.2±5.1 | 42.9±4.6; primate L2/3 |L4, p7=15.8±2.1 | 23.5±2.7, p75=50.3±4.1 | 32.7±4.9, n = ≥6m soma/dataset). Again, like excitatory connections, the surprising similarity in synaptic addition on somas across mice and primates, prompted us to check somatic synaptic density in older mice as compared to ’juvenile’ primates. At later ages, we saw our first difference in synaptic development across species. The number of inhibitory synapses on primate excitatory soma reduced in both L2/3 and L4 of V1 from ages p75 to p3000 (**Fig.3a-b, blue soma reconstructions and data points**) (mean±sem #synapses/soma for primate L2/3 |L4, p75=50.3±4.1 | 32.7±4.9, p3000=16.1±2.1 | 14.2±1.4, p = 2.366e-3(L2/3), p=4.77e-3 (L4)). Unlike primate, mouse L2/3 and L4 neurons gradually increase, or perhaps plateau, in their number of somatic synapses by the latest age we sampled (i.e., p523) **(Fig.3a-b, red somata reconstructions and data points)**. (mean±sem #synapses/soma for mouse L2/3 |L4, p105=72.2±5.1 | 42.9±4.6, p523=79.2±7.1 | 53.8±7.5, p = 0.6(L2/3), 0.1(L4)). We next reconstructed the inhibitory axons making these somatic synapses in L2/3. For both species, we found the increased soma synapse formation was the result of more axons innervating soma rather than the number of somatic synapses per axon increasing **(Fig.3c)** (mean±sem #axons/soma: mouse L2/3 p14 = 17.25±2.5, p105 = 33±2.5, p=0.016, primate L2/3 p7=13.3±2.7, p75=22.4±1.6, p=0.048, n = all soma-innervating axons across ≥ 6 soma/dataset). Likewise the number of axons innervating soma declined in primate from p75 to p3000 but remained the same, if not slightly increased, in mice from p105 to p523 (mean±sem #axons/soma: mouse L2/3 p105=33±2.5, p523=39.3±1.0, p=0.11, primate L2/3 p75=22.4±1.6, p3000=12.5 ± 1.5, p=0.013, n = all soma-innervating axons across ≥ 6 soma/dataset). These results further show that synaptic pruning on primate excitatory neuronal somatas, and not in mice, is axonal pruning (i.e., not simply pruning of synapses while maintaining the numbers of innervating axons). Indeed for primate excitatory neurons we see a dramatic ≈50% increase and reduction in the number of inputs per soma from ages p7 to p75 and p75 to p3000, respectively.

**Figure 3.**
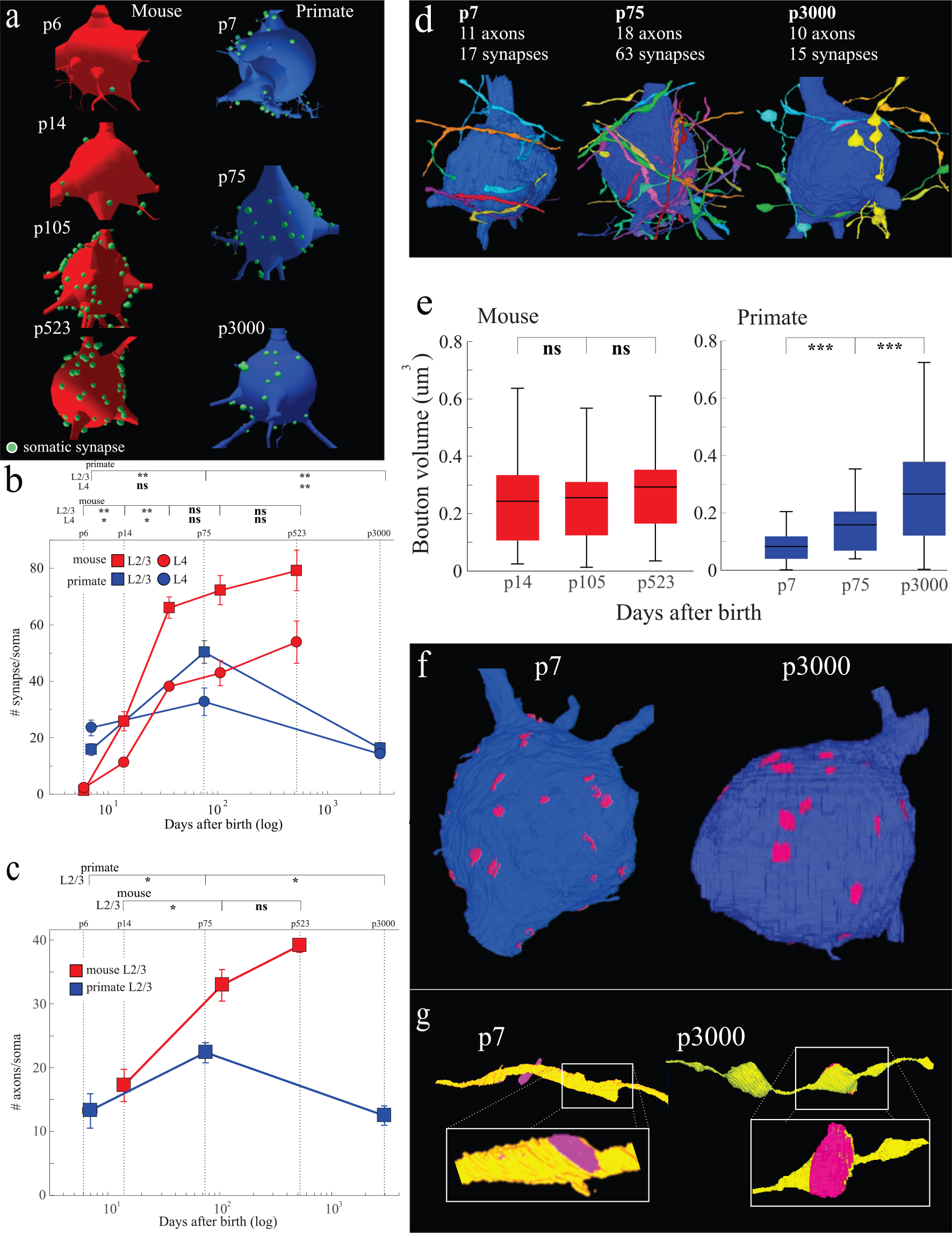
Soma innervating inhibitory axons develop differently in mouse and primate. a, Representative reconstructions of mouse (left, red) and primate (right, blue) V1, L2/3 excitatory somata at the noted postnatal ages. Green dots mark the positions of all soma synapses on each neuron. **b,** Scatter plot of x:postnatal age (log) and y:the total number of somatic synapses/soma. Blue lines = primate, red lines = mouse. Squares = L2/3, circles = L4. Smaller squares and circles display individual data points for each soma. Top line postnatal (p) days. **c,** Scatter plot of x:postnatal days (log) versus y:the total number of axons/soma for primate (blue) and mouse (red) V1, L2/3 excitatory neurons. Top line postnatal (p) days. **d,** Representative 3D reconstructions of excitatory soma and soma-innervating axons from primate V1 L23. Each reconstruction lists the postnatal age, total number of innervating axons, and total synapses found on the depicted soma. **e,** Boxplot of mouse (left red) and primate (right,blue) V1, L2/3 soma synapse bouton volume. Black lines = mean. **f,** Representative 3D reconstruction of V1, L23 excitatory neuron from p7 and p3000 primate marking PSD size (pink) of each somatic synapse. **g,** Close-up of 3D reconstruction of one soma-innervating axon from p7 and p3000 depicting the qualitative difference in bouton PSD size. Two-sample Mann-Whitney U test, ns = P > 0.05, * = P < 0.05, ** = P < 0.01, *** = P < 0.001 shown for pairwise comparisons between adjacent ages in each plot. See Supplementary Table 2 for numerical summary and supplementary files for all pairwise p-values. Lines that connect datapoints in scatter plots (b,c) are for visualization purposes only and do not represent fitted curves. Scale bar = 1 µm (e), 3 µm (g) and 300 nm (g insets). **b,** mean±sem # synapses/soma; mouse L2/3 | L4, p6 = 1.2 ± 0.3 | 2.2 ± 1.0, p14 = 25.8 ± 3.5 | 11.3 ± 1.4, p36 = 66 ± 3.8 | 38 ± 0.82, p105 = 72.2 ± 5.1 | 42.9 ± 4.6, p523 = 79.2 ± 7.1 | 53.8 ± 7.5. primate L2/3 |L4, p7 = 15.8 ± 2.1 | 23.5 ± 2.7, p75 = 50.3 ± 4.1 | 32.7 ± 4.9, p3000 = 16.1 ± 2.1 | 14.2 ± 1.4, n = ≥ 6 soma/dataset. **c,** mean±sem #axons/soma: mouse L2/3 p14 = 17.25 ± 2.5, p105 = 33 ± 2.5, p523 = 39.3 ± 1.0; primate p7 = 13.3 ± 2.7, p75 = 22.4 ± 1.6, p3000 = 12.5 ± 1.5, n = all soma-innervating axons across ≥ 4 soma/dataset. **d,** mean±sem bouton volume (µm^3^), mouse L2/3 p14 = 0.24 ± 0.03, p105 = 0.26 ± 0.02, p523 = 0.29 ± 0.03, primate L2/3 p7 = 0.08 ± 0.01, p75 = 0.16 ± 0.02, p3000 = 0.27 ± 0.03, n = ≥40 boutons/dataset.

Our discovery of inhibitory axonal pruning on excitatory somata of primate neurons **(summarized in Fig.3d)** prompted us to investigate consequences of this process. We found that after pruning, the size of the remaining boutons making somatic synapses on primate excitatory neurons grew significantly **(Fig.3e and Supplementary Fig.3)** (mean±sem bouton volume, primate L2/3 p7=0.08±0.01, p75=0.16±0.02, p3000=0.27±0.03, n = ≥40 boutons/dataset, see Supplementary Table 2 and files for all pairwise p-values). Indeed, changes in bouton volume between primate p7 and p3000 was qualitatively apparent in our 3D reconstructions when the surface area of the PSD was overlayed onto the post-synaptic soma **(Fig.3f, pink areas=PSD 3d reconstructions, and 3g, representative axon (yellow) with one depicted PSD in pink)**. The size of mice somatic synapses, which showed little sign of pruning, remained relatively unchanged over the period we examined **(Fig.3e)**. These results suggest, like in other systems with synaptic pruning [1, 61], ’surviving synapses’ become strengthened by becoming larger [67]. We also note that the total synaptic occupancy of the soma at any age was less than 10% of the total soma surface (data not shown, but see Fig.3d,f-g). Thus, it’s unlikely that pruning is driven by competition for limited physical space. Rather, decisions on which axon prunes and which do not might be more likely driven by other mechanisms such as activity. These results collectively demonstrate, for the first time, evidence of inhibitory synaptic pruning in the primate but not the mouse, indicating that the synaptic development of somatic inhibitory inputs may mark an important evolutionary distinction between these two species.

### 3.4 Saturated reconstruction and comparison between mouse neonate (p14) and adult (p105) V1

We next performed algorithmic assisted ’saturated’ reconstruction of the developing mouse brain to:

- More exhaustively analyze synaptic development in mouse cortex, given the surprise of finding little evidence of net synaptic pruning on postnatal mouse neurons.
- More quantitatively measure changes in synaptic size, map changes in non-synaptic morphologies like filopodia, and the development of sub-cellular organelles.
- Provide a resource of large volumes and annotations to the community (see Methods for data sharing plan).

Thus, we scaled and parallelized on Argonne National Laboratory supercomputers, a custom algorithmic pipeline for creating 3D EM volumes, tracing neuronal processes and identifying their connections, and incorporating human error checking **(Supplementary Fig.4)**. We used this algorithmic pipeline to analyze 91 neurons, 195 of their dendrites, and 551,652 synapses (of which 20,318 were manually proofread) on L4 mouse neurons at p14 and p105 to more thoroughly analyze the lack of pruning we observed from our manual annotations. Specifically, we utilized a combination of a flood filling network (FFN) [connected components (i.e., watershed) [for saturated segmentation of neurons and UNet + 3d, 70, 71, 72, 73] to segment synapses and mitochondria for quantifying developmental changes in synapse size, filopodia, and mitochondrial number, morphology and size. From the automatic segmentation results, we proofread 88 dendrites from the p14 dataset originating from 41 cell bodies and 107 dendrites from the p105 dataset from 50 cell bodies, overall proofreading 6,809 synapses at p14, 13,509 synapses at p105 **(Figure 4a-c)**. Overall, the cable length fully annotated of dendrites in p14 data reaches 15.4 mm and in p105 reaches 19.5 mm.

**Figure 4.**
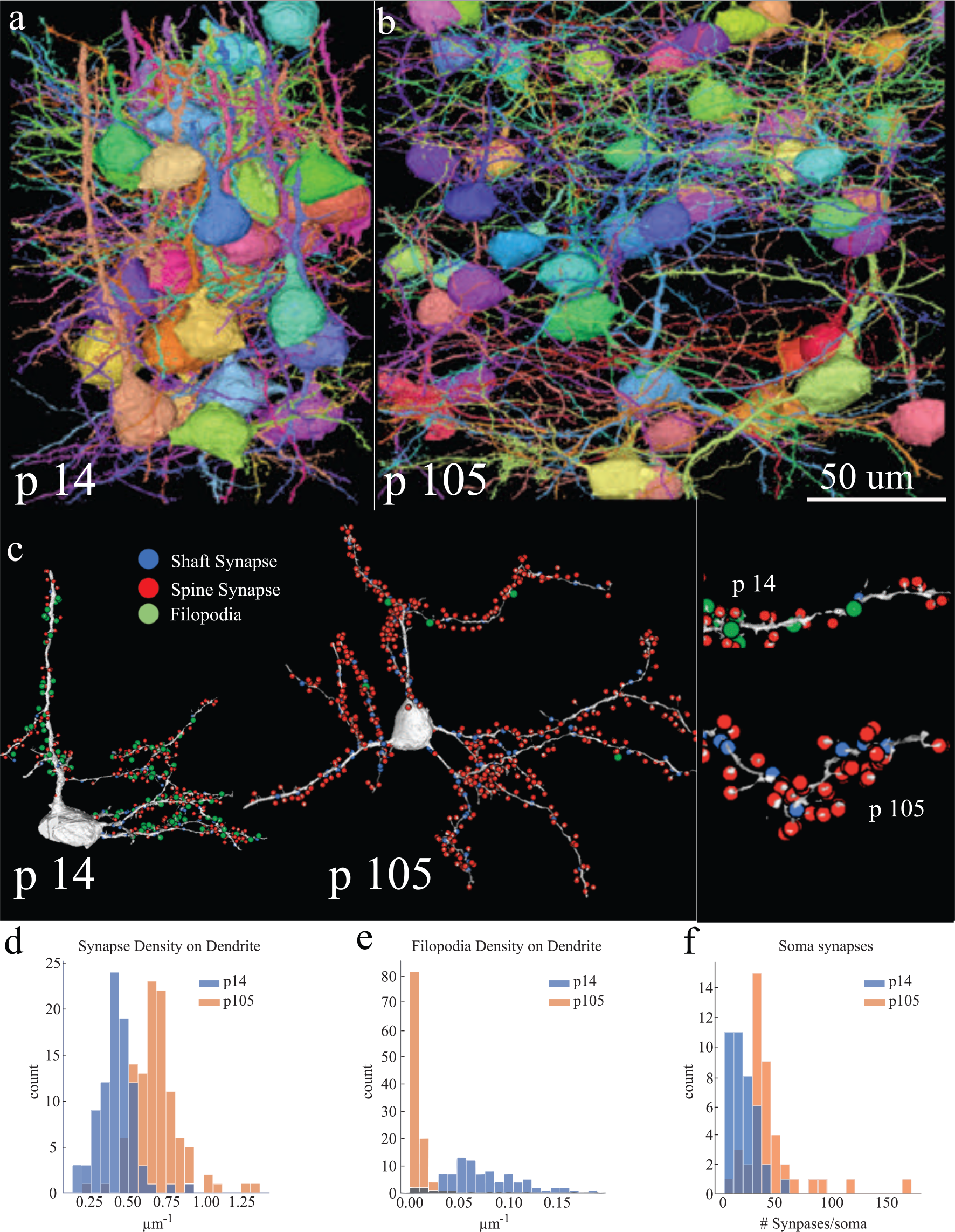
Automatic segmentation of mouse V1 p14 and p105 excitatory neurons. a-b, Representative reconstructions of L4 excitatory neurons from mouse V1, p14, p105. Each neuron is uniquely colored for visual purposes. **c,** Representative reconstruction of a single mouse V1 p14 and p105 excitatory neuron with the position of inhibitory shaft synapses (blue dots), excitatory spine synapses (red dots), and filopodia (green dots) detected using automatic segmentation. *Right*, zoom in of left images showing a dendrite from p14 (top) and p105 (bottom). **d-f,** Histograms of mouse V1 p14 (blue) and p105 (orange) comparing distribution in the frequency of (**d**) excitatory spine synapses/µm, (**e**) filopodia/µm, and (**f**) total number of inhibitory synapses/soma. Two-sample Mann-Whitney U test. Scale bar = 25 µm (a). **d,** mean±sem spine synapses/µm; p14 = 0.43 ± 0.11, n= 88 dendrites, p105 = 0.69 ± 0.16 synapses/µm, n = 107 dendrites, p = 6.17e-26. **e,** mean±sem filopodia/µm; p14 = 0.075 ± 0.038, n = 88 dendrites; p105 = 0.005 ± 0.008, n = 107 dendrites, p = 4.43e-32. **f,** mean±sem total synapses/soma; p14 = 18.31 ± 12.15, n = 39 soma; p105 = 42.85 ± 28.80, n = 41 soma, p=2.65e-8.

#### 3.4.1 Further evidence of primarily synapse formation in developing mouse visual cortex

We observed a mean±sem synapse density of 0.43±0.11 synapses/µm across 88 dendrites at p14 and 0.69±0.16 synapses/µm across 107 dendrites at p105, amounting to a 60.4% increase (p=6.17e-26) **(Fig.4d)**. We believe the differences from the automated segmentation data and our hand tracings are likely due to an over-representation of large proximal dendrites attached to soma in our automated analyses (see Sup. Figure 2). We also manually annotated filopodia along these dendrites and found a significantly higher density in p14 data, while in p105, filopodia are extremely rare (mean±sem: p14=0.075±0.038, p105=0.005±0.008, p=4.43e-32) **(Fig.4e)**. We used the distinct morphological criteria of filopodia as long protrusions from dendrite that do not form a postsynaptic structure [74, 75]. Finally, we also found a similar rise in the number of somatic synapses from p14 to p105 (mean±sem: p14=18.31±12.15, p105=42.85±28.80, n = 39 and 41 soma, respectively, p=2.65e-8) **(Fig.4f)**. Overall, these results from automatic segmentation are consistent with and provide further validation of our manual annotation.

#### 3.4.2 Increase in synapse size

A major advantage of connectomics is its ability to resolve ultra-structural neural morphology. With a combination of automatic mask prediction and manual correction, we were able to quantitatively measure each synapse for its vesicle size and PSD size **(Supp Fig.5)**. We compared the distribution of these metrics at p14 and p105 and observed a significant increase in both synaptic junction size (66.5%) and in vesicle cloud size (14.8%) in p105 mice (both p < 1e-5) **(Supp Fig.6)**, suggesting an overall maturation of synapses. Interestingly, we find that synapse size distributions are primarily log-normal, which has also been observed in synapse size distributions in the adult mouse brain [76, 77], suggesting that such distributions are not the result of developmental synapse re-arrangements. Thus, we conclude that the primary development of synapses in mouse primary visual cortex is the addition of new synapses and their growth with the concurrent removal of large numbers of filopodia. Finally, we find that the average distance of a spine synapse from ’parent’ dendrites is similar at both p14 and p105 **(Supp Fig.7)** (mean±sem distance to dendritic branch (nm); p14 = 1107 ± 9.7, p105 = 1081 ± 7.5, p = 0.80) despite a 60.4% increase in the number of synapses, suggesting intrinsic limitations to the distances spines can extend, independent of the number of synapses formed and unaffected by the process of development.

#### 3.4.3 Mitochondria size development and correlation with synapse density

Our segmentation pipeline allows us to reconstruct sub-cellular organelles in addition to synapses, allowing the development of organelles to be correlated with the development of synapses. As an example, we addressed how mitochondria develop and how that development correlates with the development of synapses. We chose mitochondria as they have been implicated in numerous synaptic functions including sustaining long-term plasticity [78, 79, 80]. We reconstructed 398,278 instances of mitochondria in p14 and 533,019 in p105. We analyzed mitochondria primarily in the dendrites of excitatory neurons and found a 78.4% increase in mitochondria size and 66.7% increase in mitochondria density (i.e., total mitochondria volume/length of dendrite analyzed) in p105 relative to p14 **(Supp Fig.8-9)** (mean±sem mitochondria size (µm^3)^; p14 = 0.04263±0.0, n = 39,8278, p105 = 0.0687 ± 1.9e-4, n=533,019, p ≈ 0; mean±sem mitochodria density (nm^2)^; p14=19,236.2 ± 571.1, n = 88, p105 = 32,072.9 ± 2,581.3, n = 107, p ≈ 1e-20). Finally, previous connectomic analyses [81] have demonstrated a positive correlation between synapse density and mitochondria coverage in basal dendrites of neurons in the mouse V1, L2/3 [81]. We found, indeed, that this correlation emerges with developmental age at age p105, there is a stronger correlation between mitochondria coverage and synapse density in basal dendrites relative to p14, but little correlation in apical dendrites in both ages **(Supp Fig.10)** (Pearson correlation coefficient, apical and basal combined: p14 r=0.12, p=0.25, p105 r=0.5,p=5.8e-8; apical: p14 r=-0.01, p=0.97, p105 r=0.18, p=0.44; basal: p14 r=0.2, p=0.1, p105 r=0.54,p=6.4e-8).

## 4 DISCUSSION

### 4.1 The clock of synaptic development is uncoupled from lifespan

We investigated the synaptic development of cortex in two foundational models in neuroscience, *Mus musculus* (mouse) and *Macaca mulatta* (primate). Given the disparate lifespans of these species, and current models that emphasize the neotonous nature of primate development[82], particularly human, [83], we specifically asked whether mouse neurons showed an accelerated cycle of synaptic formation and elimination relative to primates.

Instead of accelerations, we saw two different developmental strategies across species based on the type of synapse. First, the development of excitatory synapses was isochronic between mice and primates (Fig. 1b), with little sign of net pruning during early mouse postnatal life, as might have been expected from the end of ’critical periods’ at this same time [84, 85, 25]. Indeed, previous results have demonstrated isochronic excitatory synaptic development across primate cortical areas in the same animal, with a time course of addition and pruning similar to the time courses observed here, [7]. Thus, we extend the idea that there is a single temporal program for excitatory synaptic development in cortex, independent of cortical area, layer, or even species, despite the surprising fact that these neurons across cortical layers are likely generated at different embryonic times [86, 87, 88] and receive synaptic connections from nearby and distal sources (e.g., thalamus) at different rates [89, 90].

Second, inhibitory synaptic development was also not accelerated in mice. However, unlike excitatory synaptic connections, the basic developmental program of inhibitory connections differed across species. Inhibitory innervation of primate excitatory neurons showed a cycle of formation and pruning similar to excitatory synaptic development but the same class of inhibitory synapses on mouse excitatory neurons showed only synaptic addition throughout life (Fig.3b-c). The striking similarity in the development of excitatory synaptic connections contrasted with different developmental programs of inhibitory synapses, suggests that changes in the nature of inhibition, either from the addition of new cell types, or as demonstrated here, changes in developmental programs, could be a source of evolutionary innovation from rodents to primates. Furthermore, our results add a new dimension in support of a growing body of genetic evidence for evolutionary novelties in inhibitory circuits [91, 92, 93, 94, 95].

We conclude that neither the development of excitatory nor inhibitory synaptic connections showed signs of classic ’heterochrony’ [48]. We argue that since such rules are based on comparisons of rates of cell division and differentiation, they are inadequate to describe the ontogeny and phylogeny of synapse development on post-mitotic mammalian central neurons. Our conclusions outline the foundation for a new model of neonatal synapse development and necessitates further investigations for new principles of synaptic development across species **(Supp Fig.11)**. It is noteworthy that we measured only net changes in synapses, not specific counts of additions and subtractions, so we cannot speak to dynamic changes (i.e., synapses that are pruned or reformed), or if a smaller population of axons are undergoing large scale synaptic re-organizations. These limitations do not undermine our conclusions but rather motivate investigations that would further refine our new model for synaptic development.

### 4.2 Relevance to Prior Literature

#### 4.2.1 Prior Reports of Cortical Synapses Pruning in Early Mouse Postnatal Life

These results favor a model of excitatory synapse formation in cortex where filopodia mature into spine synapses [56, 96, 74, 97, 98, 75, 99] as opposed to other models including transformation of shaft synapses into spines, or spine formation followed by pruning **(Supp Fig.12)**. The widespread prevalence of filopodia in p7-p14 neurons **(Fig. 4e)**, which can be hard to differentiate from spines without unambiguous identification of a pre-synaptic axonal partner, could have potentially inflated counts of synaptic density in reports using lower resolution and sparse labeling optical microscopy. There are also numerous reports of changes, potentially heterochronic changes, in gene transcription [83, 100, 101], sometimes even of genes implicated in synapse formation and pruning [102]. However, since we are uncertain about the ’conversion’ factor (i.e., how many additional RNA transcripts correlate to an additional synapse), we remain uncertain how to relate our findings to those. Synaptic pruning of inhibitory synapses in the first few weeks of postnatal life in mice has been suggested by numerous studies [103, 37, 104]. Inhibitory synapses are harder to investigate by light microscopic investigation (e.g., no spines as a proxy for connectivity). Thus, evidence for developmental changes in inhibitory circuits relies even more on indirect measures of connectivity including pharmacological intervention [105], immuno-histochemistry [106], electrophysiological characterization [107], putative trans-synaptic labeling [28, 29, 30], genetic manipulations [108], and transcriptional changes [109]. We provide the first complete accounting of every somatic, putatively parvalbumin inhibitory synapse, across species and find there is no net pruning in inhibitory connections over mouse postnatal life, extending previous results that used sparse sampling EM without identification of somatic vs. shaft inhibitory synapses [37]. Our results call for a revaluation of the role of inhibition on critical periods and the effect of experience on inhibitory circuity [110, 111, 112].

Finally, there is a broad consensus with our primary result: a lack of net excitatory synapse pruning in the first few months of postnatal life across species covering 4 phylogenetic orders (Supp Fig.13) (see Methods) [9, 10, 11, 12]. Our results provide a potential cautionary tale for comparing mouse and non-human primate brains across developmental time points. For example, we previously reported sparse innervation of ’adult’ primate neurons as compared to ’adult’ mouse neurons [38]. Our work here suggests instead that we may have been comparing different parts of an similar curve of synaptic development. We argue that potentially similar confusions could occur in mouse models of human neurological disease, particularly models assuming synaptic overabundance and pruning in early mouse postnatal cortex (e.g., mouse models of Autism Spectrum Disorders [20, 21, 22]). We argue that large volume EM connectomes of such disease models serves as a complete, unbiased, and standard reference to evaluate connectivity changes across models and interventions and aid in translating findings in mice to humans.

#### 4.2.2 Conservation of the Mechanisms of Synaptic Pruning

We find several noteworthy and novel demonstrations of cortical synaptic pruning. First, our analysis of excitatory axons reveals that cortical synaptic pruning during mouse and primate development follows similar cellular mechanisms as reported from a variety of peripheral systems, particularly the mouse neuromuscular junction (NMJ). Like the NMJ [1, 61], dendrites of primate excitatory neurons shows sparse anatomical inputs at early ages, a dramatic increase in the number of axonal inputs followed by axonal/synaptic pruning. As in the NMJ, we find, during periods of net synaptic pruning, numerous examples of mitochondria-filled ’axonal retraction’ bulbs (Fig.2d-f), an anatomical signature of axonal pruning at NMJs. These similarities suggest that axonal retraction might be a common mechanism for synaptic rearrangement in the nervous system. Interestingly, our previous observations that such retraction bulbs are formed during exposure to drugs of abuse [113] suggests that this common mechanism might also extend to pathological conditions of brains. Second, our analysis of inhibitory axon pruning in primate reveals an interesting principle in how axons rearrange during development. Somatic synapses that survive developmental pruning increase in size (Fig.3e-g). Synapse size, specifically, the size of the PSD has a linear correlation with the amplitude of the excitatory postsynaptic potentials (EPSPs) [67], suggesting the somatic synapses that survive pruning are also stronger. Future investigations will help determine the mechanisms of pruning of inhibitory synapses in primate cortex (i.e., does pruning select previously strong synapses or does strengthening emerge from the process of pruning).

## Supporting information

pairwise p-values

supplemental species data

## Acknowledgements

Primate p7 and p75 tissue was kindly provided to us by Dr. Alvaro Duque of the MacBrain Resource Center (MBRC) of Yale School of Medicine. The MacBrain Resource Center is supported by NIH grant R01MH113257 (to Dr. Alvaro Duque). We would like to sincerely thank Vandana Sampathkumar, Dawn Paukner, and Anastasia Sorokina for helping with manual proofreading of the automatic segmentation. We would also like to thank Dr. Kevin M Boergens, Professor Murray S. Sherman, and Professor John Maunsell for their thoughtful suggestions on the manuscript preparation.

## 5 Methods

### 5.1 Animal subject and tissue acquisition details

Brain tissue processed in our lab (i.e., mouse V1 p6,p14,p105,p523 and primate V1 p7,p75,p3000) was prepared for EM as previously described [114]. Dissection of V1 and serial electron microscopy imaging was performed as previously described [38, 115]. Primate p7 and p75 tissue was kindly provided to us by Dr. Alvaro Duque of the MacBrain Resource Center (MBRC) of Yale School of Medicine. The MacBrain Resource Center is supported by NIH grant R01MH113257 (to Dr. Alvaro Duque).

### 5.2 Data collected

Total volumes imaged and links to external datasets are listed in Supplementary Table 3.

### 5.3 data availability

Homemade code used for EM image processing (i.e., 2D stitching, 3D alignments, brightness/contrast normalization) can be freely accessed here: . 3D alignment was performed using the publicly available Aligntk software available here: https://mmbios. pitt.edu/aligntk-home. Homemade code used for data analysis of Knossos traced skeletons (i.e., xml files) can be freely accessed here: nal EM data for V1 are freely available online at bossdb.org. Mouse S1 datasets that we analyzed for changes in excitatory synapses are available through original publication [40]. Adult S1 excitatory synapse measurements were manually analyzed using the publicly available neuroglancer file from [115] which can be found here: https://github.com/google/neuroglancer; Kasthuri et al., 2014. Mouse somatosensory cortex(6x6x30 cubic nanometer resolution). Mouse p36 datasets are publicly available here for L2/3: and here for L4: https://www.microns-explorer.org/cortical-mm3. Automatic segmentation of neurons, synapses and mitochondria of mouse V1 L4 p14 and p105 are available as a WebKnossos format available for further public proofreading/error checking.

### 5.4 Manual segmentation

Data annotations were done using Knossos (https://knossos.app/) and performed by 3 individuals (GW, HL, BK). To ensure accuracy of the data, 33% of the annotations from one person was verified by the other. 33% of the annotations were given to a naïve annotator to verify the accuracy. We found a >98% agreement between manual annotators. We found that 100% of spines could be reconstructed. Classes of neurons and their dendrites were identified by distinguishing anatomical properties [38]. We used the following metrics to identify and quantify each anatomical feature reported on: 1. Synapses were identified by the presence of a post-synaptic density and vesicles on the pre-synaptic axon. 2. Spine and shaft synapse frequency: a dendrite was chosen at random and traced for 10 µm to first determine if it was an excitatory dendrite. The number of spine or shaft synapses contained within the 10 µm window were counted manually in Knossos and the total number of synapses were divided by the actual segment length to calculate spine synapses/µm and shaft synapses/µm. The diameter of the dendrite was calculated by measuring across the diameter in all three orthogonal views and then averaged. 3. Soma synapses: excitatory soma were identified as being apart of neurons with spinous dendrites. Neurons whose soma was fully within the imaged volume were used to count the total number of soma synapses. Perisomatic synapses were scored along the first 10µm of dendrite that left the soma. 4. Bouton size: a node (i.e., sphere) was placed over a bouton in knossos and sized to best fit the size of the bouton. The node radius was used to calculate the surface area using S.A = π4r2

### 5.5 3D rendering

All 3D manual segmentation rendering was done using VAST [116] or neuroglancer.

### 5.6 Statistical methods

Quantification and statistical tests of anatomical measurements Mean and standard error of the mean (SEM) was calculated for every quantification. Statistical significance was calculated using the two-tailed Mann-Whitney U test [117] between aggregate mouse and aggregate primate datasets. Mann whitney U implemented in r using wilcox.test. See Table 1-2 and Figure Legends for the exact value of (n) and what (n) represents. All pairwise p-values available in excel format in Supplementary files. Box plots indicate variability outside the upper and lower quartiles and center black line indicates the mean.

### 5.7 Automatic segmentation

An extensive description of all automatic segmentation detail can be found here:

### 5.8 scraped data from the existing literature

To generate Supplementary Figure 13, we first conducted an extensive literature search to collate publications that reported synapse density measurements across development in different species. We eliminated those that were missing early ages or those that used indirect methods (e.g., golgi labeling and spine counting). After these filters, we identified four papers [9, 10, 11, 12]. For a paper that listed synapse density values in a table, we used those values directly. For papers that showed results as a line/scatter plot rather than a table, we extracted values from the plot for ages that span the animals life starting with p7 as the earliest data point. Each dataset was then normalized to a non-dimensional value by first setting p7 to zero and dividing all other postnatal time point value by the p7 value to calculate a % increase relative to the p7 value (e.g, if p7 = 250 and p15 = 2500 synapses/neuron, p7 set to 0 and p15 to (2500/250)=10. See supplementary excel file ”species synapse scrape” for full normalization of each dataset.

## SUPPLEMENTAL FIGURES

**Supplementary Figure 1.**
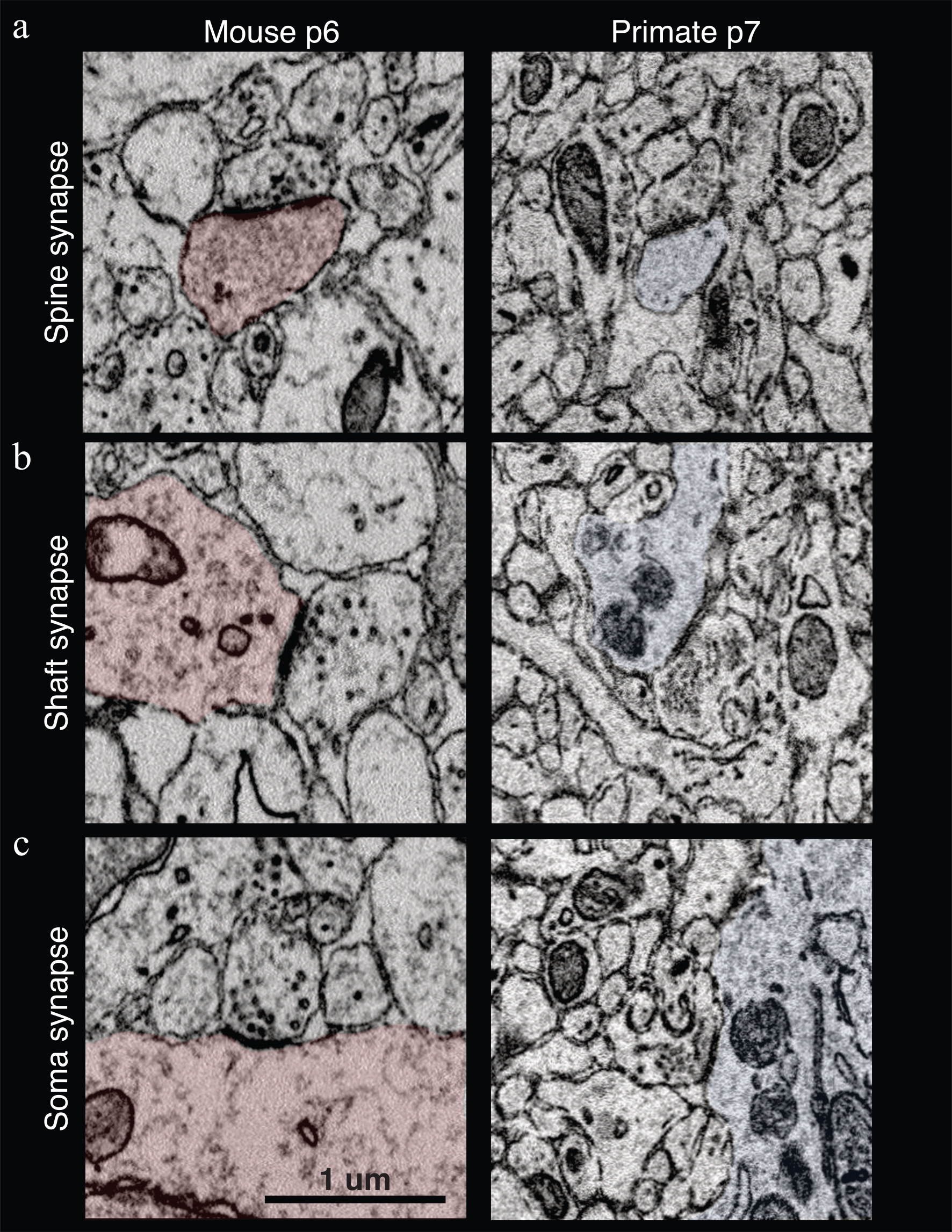
Synapses of the p6 mouse and p7 primate. Representative 2D EM images of **(a)** excitatory spine, **(b)** inhibitory shaft, and **(c)** inhibitory soma synapses in p6 mouse (left) and p7 primate (right). In each image, the post-synaptic target is colored in red (mouse) or blue (primate). Note the prominent PSD and pre-synaptic vesicle cloud established early in neonatal development. Scale bar = 1 µm.

**Supplementary Figure 2.**
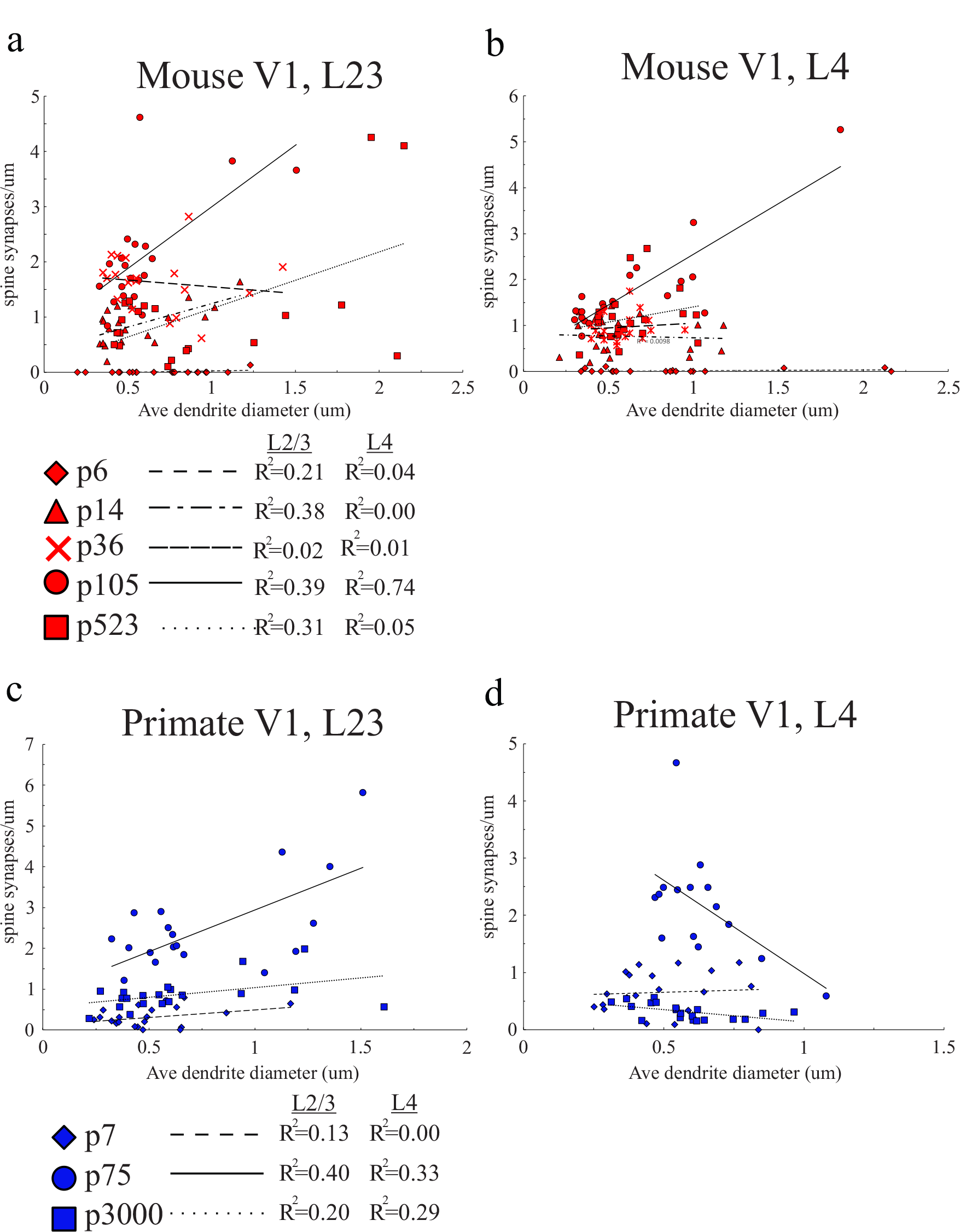
Excitatory spine synapse density relationship to dendrite diameter. Scatter plot of x: average diameter of quantified 10 µm dendrite fragment and y: spine synapse/µm along the 10 µm dendrite fragment for mouse V1, L2/3 **(a)** and L4 **(b)**, and primate V1, L2/3 **(c)** and L4 **(d)**. Key for mouse (red) and primate (blue) show line-type representing best linear fit along with corresponding r^2^ value for each data set. Linear regression used for r^2^ calculation.

**Supplementary Figure 3.**
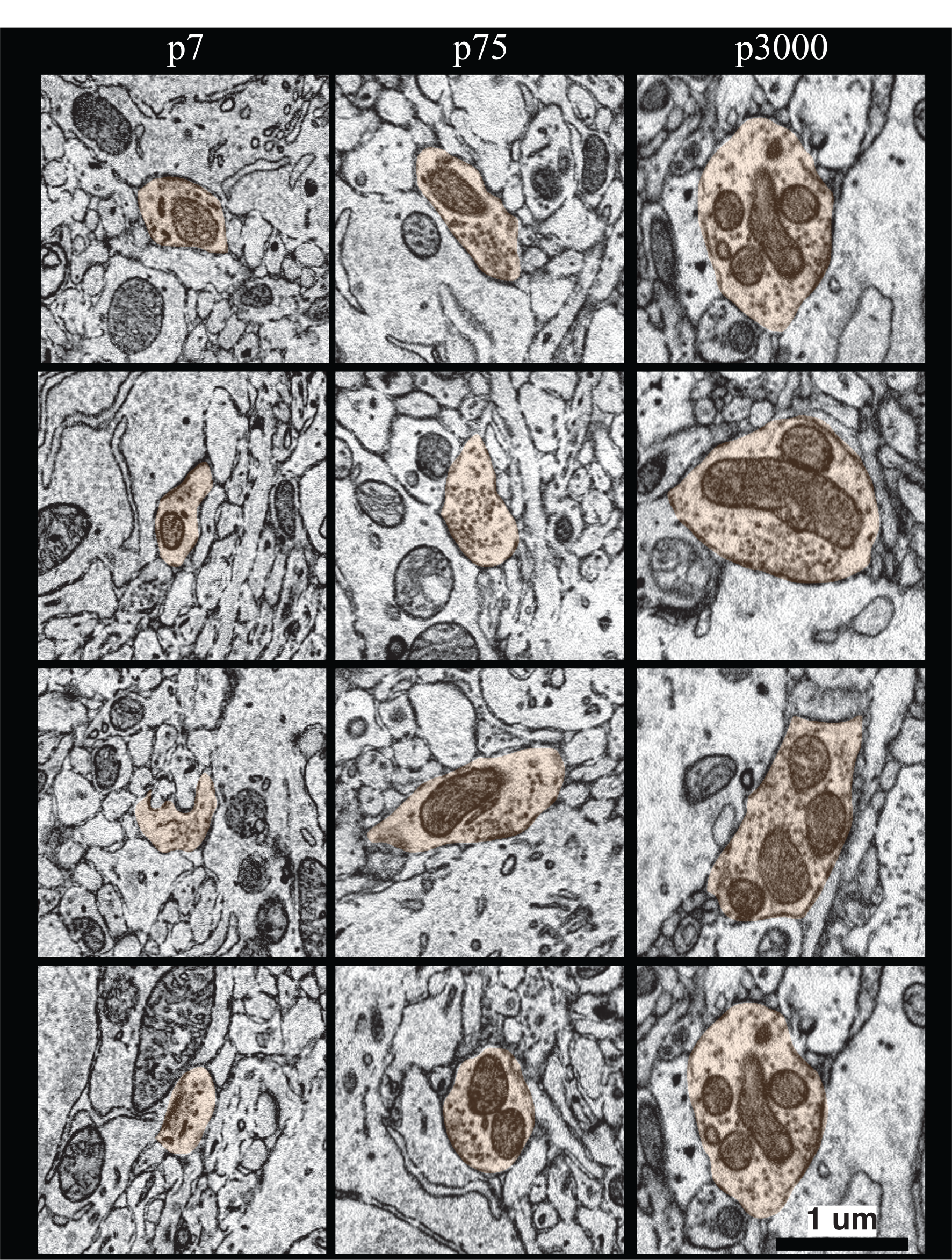
Boutons of synapses onto primate excitatory neuron soma increase in size over development. 2D EM images showing a column of four representative axonal boutons (highlighted in orange) making a synapse onto primate excitatory axons at the noted ages. Scale bar = 1 µm.

**Supplementary Figure 4.**
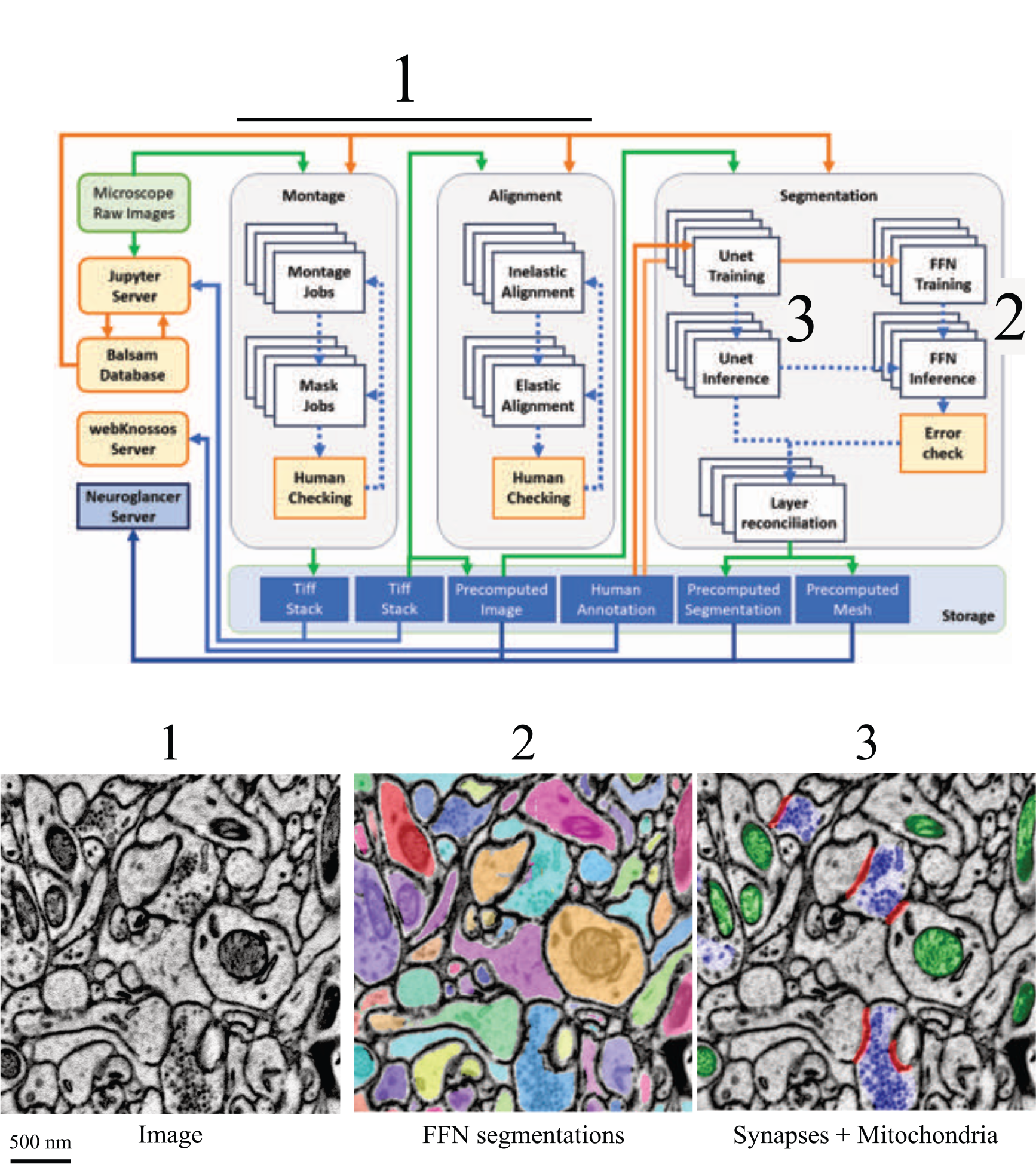
Schematic of the automatic segmentation pipeline. *Top:* The green box represents the data acquisition. Orange boxes and arrows represent human interactions in the pipeline. The white boxes represent High Performance Computing (HPC)-submitted jobs. Blue boxes represent the data storage and visualization server. Green arrows represent I/O from the computing resources. *Bottom:* Representative images generated from different steps in the pipeline: 1 = raw images produced from 2D montage and 3D alignment, 2 = FFN saturated segmentation of all neuronal processes, and 3 = UNET/connected components sparse segmentation of synapses (red=PSD and blue=vesicle cloud) and mitochondria (green). Scale bar = 500nm.

**Supplementary Figure 5.**
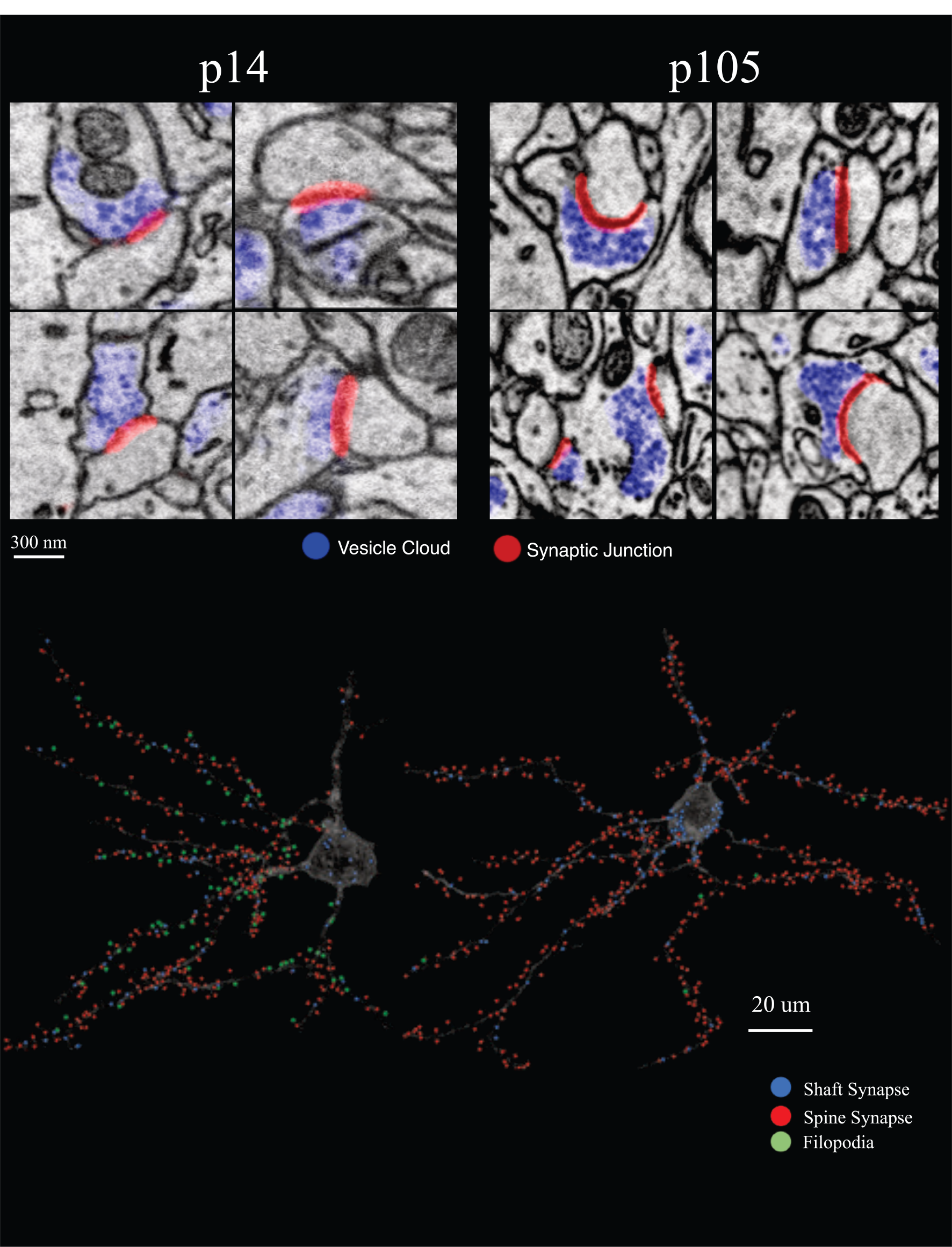
Automatic segmentation of synapse vesicle cloud and PSD. *Top:* Representative 2D EM images showing automatically identified synapses in mouse p14 and p105 datasets. blue colored regions mark the pre-synaptic vesicle cloud, and red lines mark the PSD. *Bottom:* Representative rendering a p14 (left) and p105 (right) neuron derived from FFN segmentation overlayed with UNET segmentation of shaft synapses (blue circles), spine synapses (red circles), and filopodia (green circles).

**Supplementary Figure 6.**
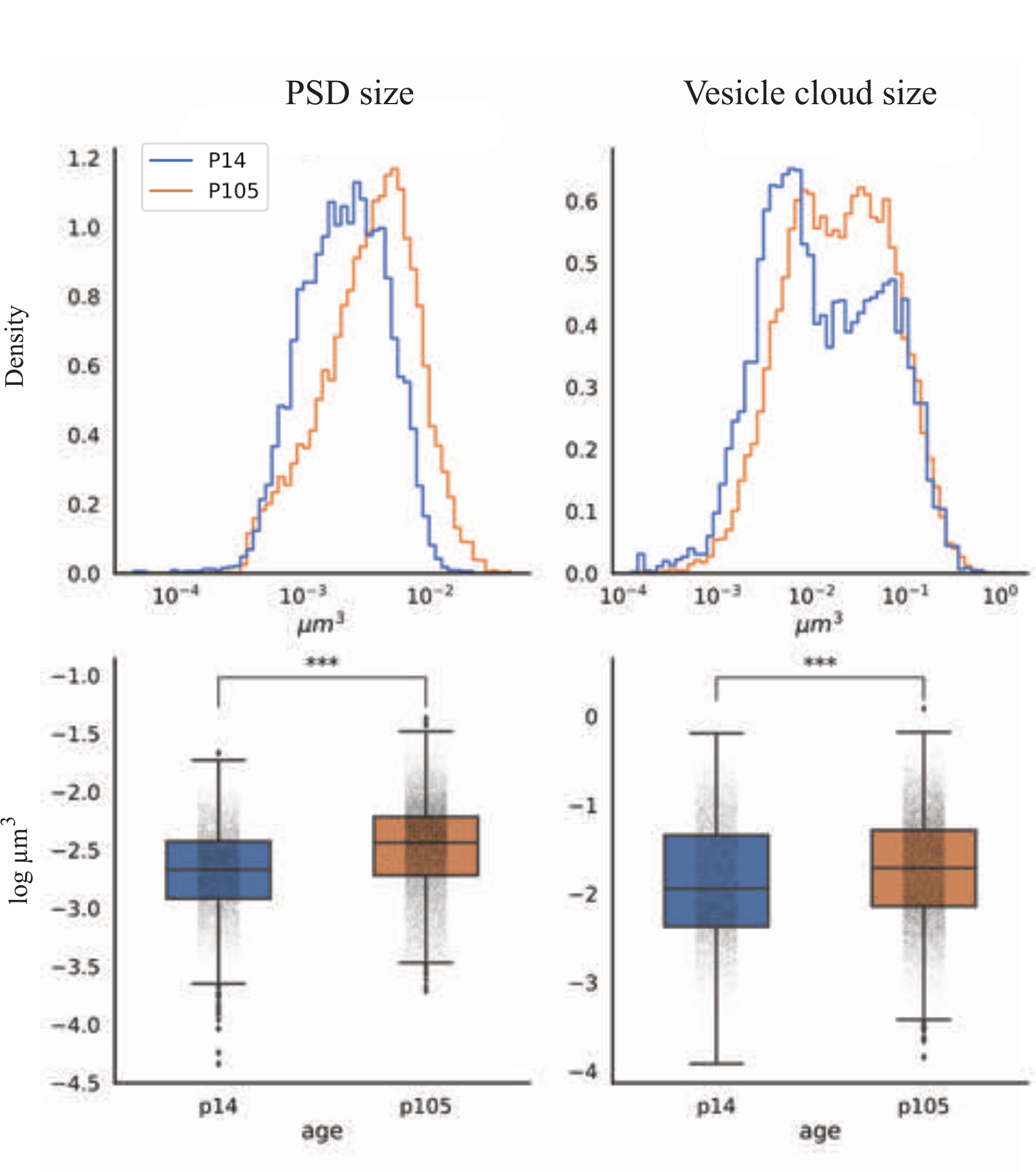
Quantification of spine synapse changes from manually-error checked, automatic segmentation in mouse p14 and p105, L4. Changes in PSD size *(Left)*, and vesicle cloud size *(Right)* between mouse p14 (blue) and p105 (orange) V1,L4. Each analysis is represented as both a density (top) and box plot (bottom). mean±sem PSD size (µm^3^); mouse p14 = 0.0028 ± 2.6e-5, n = 6732, p105 = 0.0046 ± 3.3e-5, n = 13478. mean±sem vesicle cloud size (µm^3^); mouse p14 = 0.035 ± 6.4e-4, n = 6780, p105 = 0.0403 ± 4.8e-4, n = 13500, p < 0.0001 for both results. Two-sample Mann-Whitney U test.

**Supplementary Figure 7.**
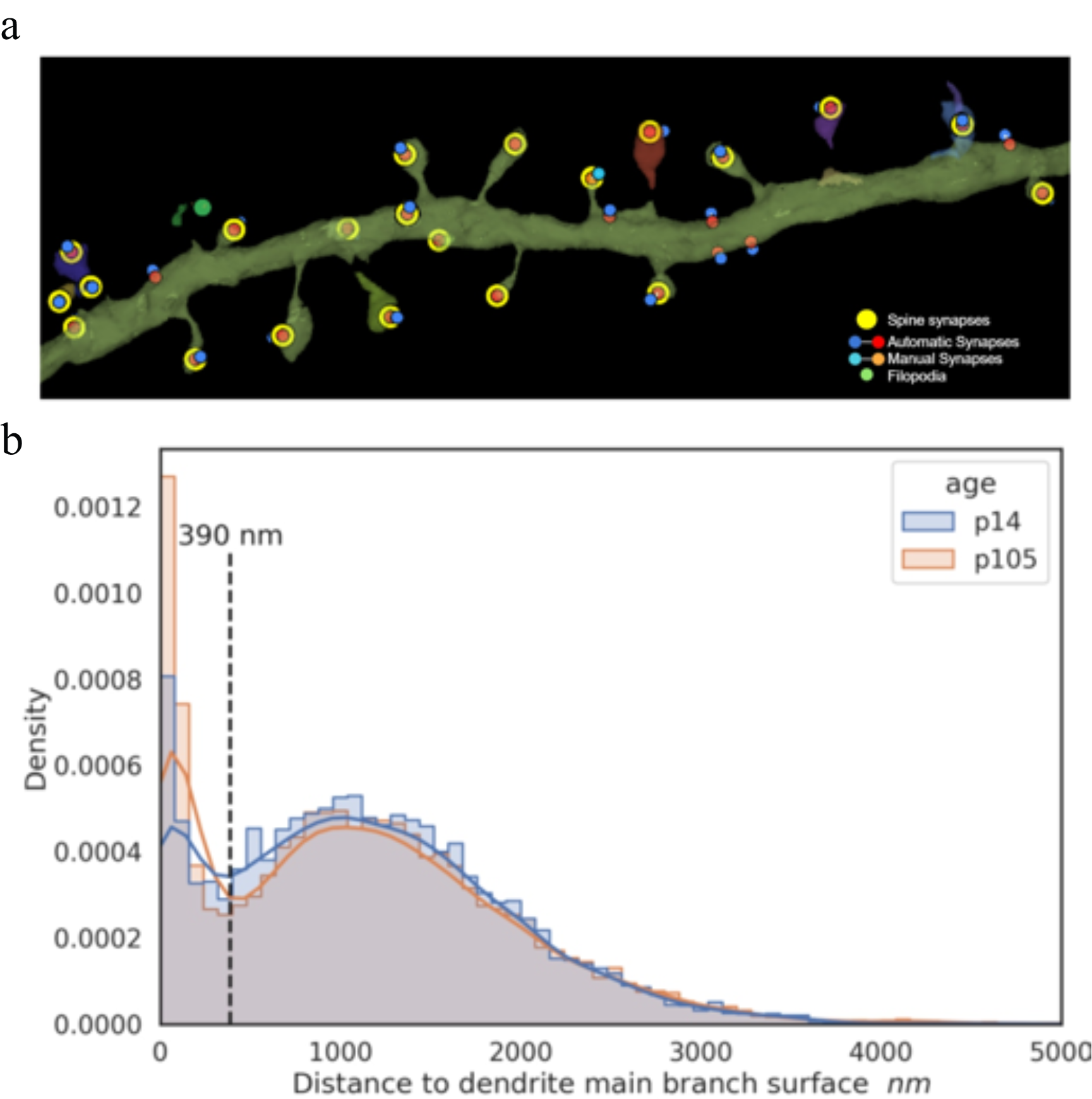
Distance of spine synapse PSD from dendritic shaft in p14 and p105 mouse V1, L4. a, Representative reconstruction of an excitatory dendrite with spine synapses marked in yellow, filopodia in green, and stick-and-ball segments from manual (dark blue-red) and automatic (light blue-orange) annotations used for measuring the euclidian distance between the PSD and dendrite shaft. **b,** Histogram of distance from PSD to dendrite surface (i.e., shaft) for mouse V1, L4 excitatory neurons in at ages p14 (blue) and p105 (orange). A 390 nm cut-off was used to differentiate shaft (<390nm) from spine (>390nm) synapses. Mean±sem distance to dendrite surface (nm); p14 = 1107 ± 9.7, p105 = 1081 ± 7.5, p = 0.80. Two-sample Mann-Whitney U test.

**Supplementary Figure 8.**
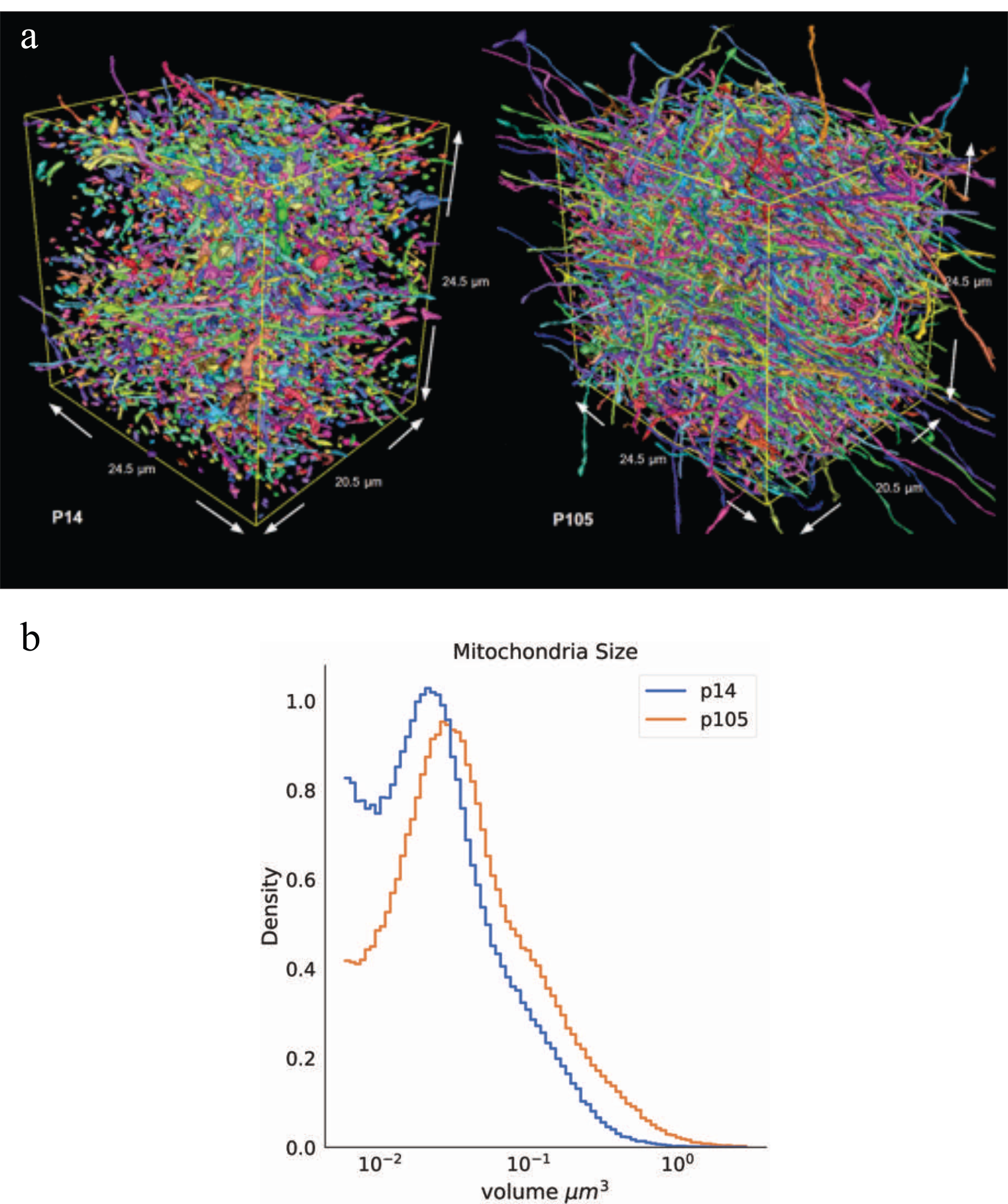
UNET automatic segmentation and analysis of changes in mitochondria size in mouse p14 and p105, L4. a, 3D rendering of a representative subvolume depicting automatically segmented mitochondria from mouse p14 (left) and p105 (right). Each colored object represents an individual mitochondria. **b,** Density plot quantifying mitochondria size from automatic segmentation in mouse p14 (blue) and p105 (orange). mean±sem mitochondria size (µm^3^); mouse p14 = 0.043 ± 0.0,n = 39,8278, p105 = 0.069 ± 1.9e-4, n=533,019, p ≈ 0. Two-sample Mann-Whitney U test.

**Supplementary Figure 9.**
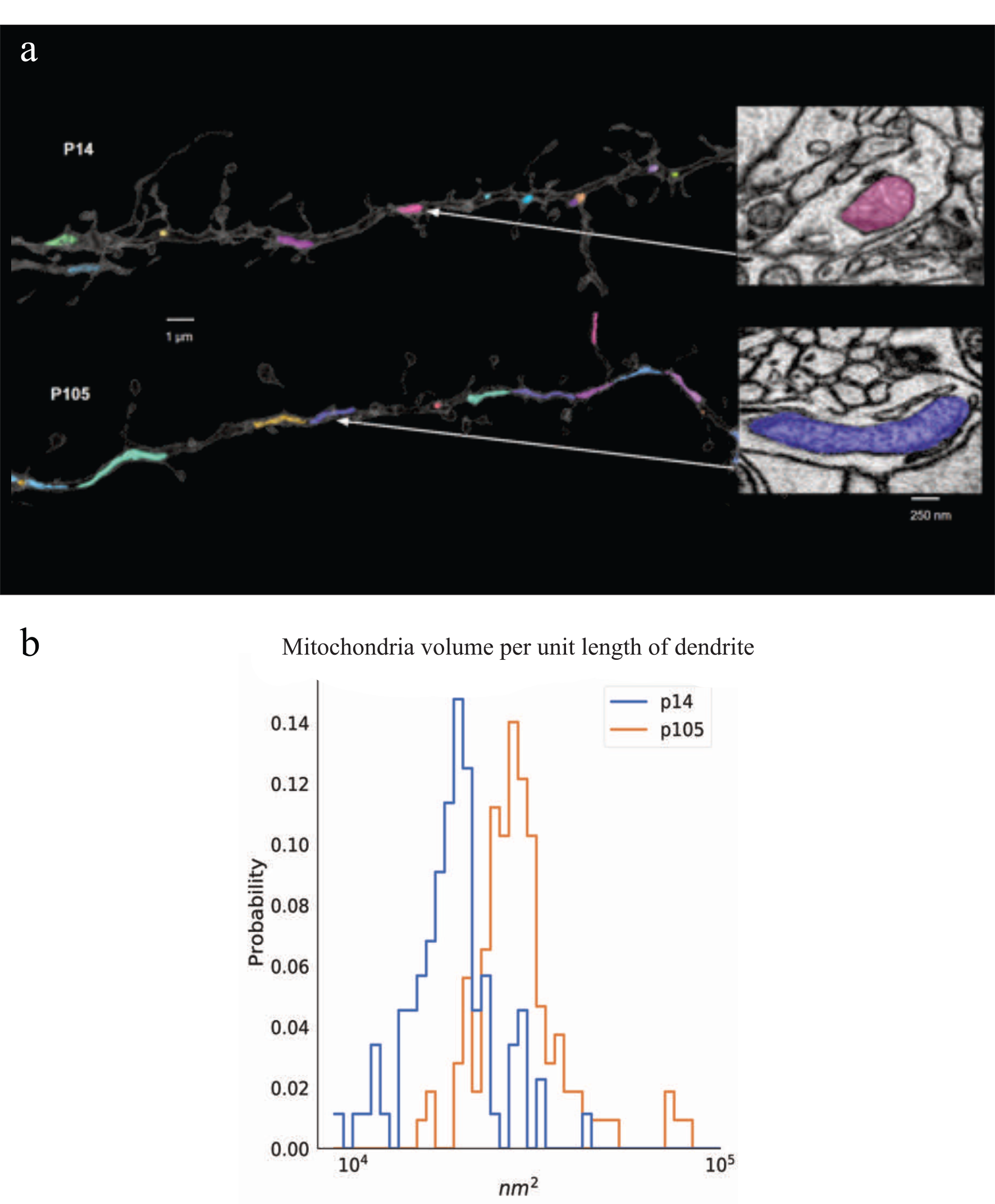
Mitochondria density along dendrites in mouse p14 and p105, L4. a, Representative rendering from FFN segmentation of an excitatory dendrite from p14 (top) and p105 (bottom) containing UNET segmented mitochondria. *Right:* 2D EM image showing an example of one segmented mitochondria from each dendrite. **b,** Density plot quantifying mitochondria volume per unit length of dendrite. mean±sem mitochondria per unit length of dendrite (nm^2^); mouse p14 = 19,236.2 ± 571.1, n = 88, p105 = 32,072.9 ± 2,581.3, n = 107, p ≈ 1e-20. Scale bar: a, left= 1 µm; a, right=200 nm. Two-sample Mann-Whitney U test.

**Supplementary Figure 10.**
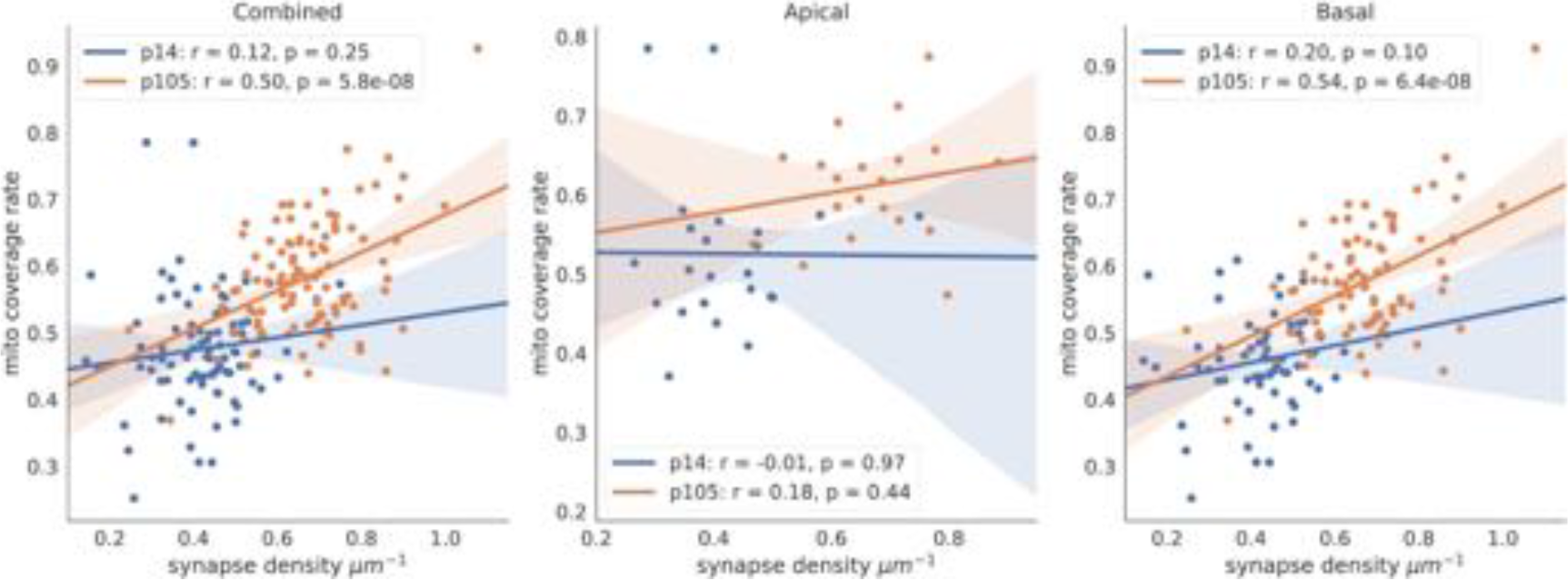
Correlation between synapse density and mitochondria coverage. Scatter plots of x:synapse density and y:mitochondria coverage rate in p14 (blue) and p105 (orange) across *left*: apical and basal dendrites combined, *middle*: apical dendrites only, and *right*: basal dendrites only. Pearson correlation coefficient, combined: p14 r = 0.12, p = 0.25, p105 r = 0.5, p = 5.8e-8; apical: p14 r = -0.01, p = 0.97, p105 r = 0.18, p = 0.44; basal: p14 r = 0.2, p = 0.1, p105 r = 0.54, p = 6.4e-8. Two-sample Mann-Whitney U test.

**Supplementary Figure 11.**
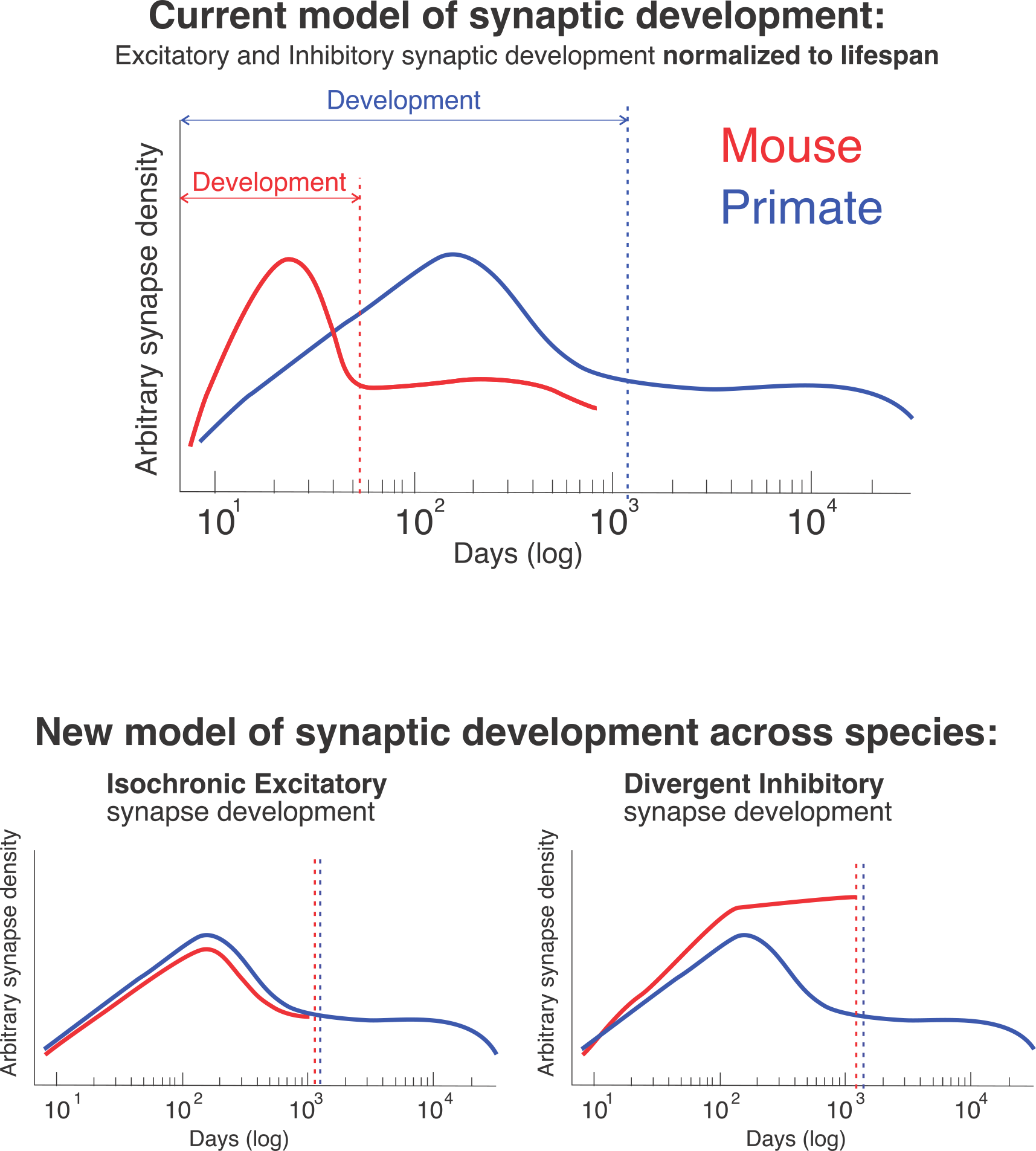
New model of excitatory and inhibitory synapse development. *Top*: Current model of synapse development in mouse and primate. The cycle of formation and pruning of both excitatory and inhibitory synapses is normalized to the animals lifespan where the cycle is compressed, in absolute time, in the mouse relative to the primate such that the cycle occurs during neonatal development in both species. *Bottom*: New model of synaptic development in mouse and primate. Excitatory synapse development is isochronic across species (i.e., a single synaptic clock) (left), whereas inhibitory somatic synapses differ across mouse and primate (right): in mouse, somatic synapses are continuously added throughout the life of the animal (red line) but in primate, somatic synapses accumulate and are pruned at a similar time as excitatory synapses.

**Supplementary Figure 12.**
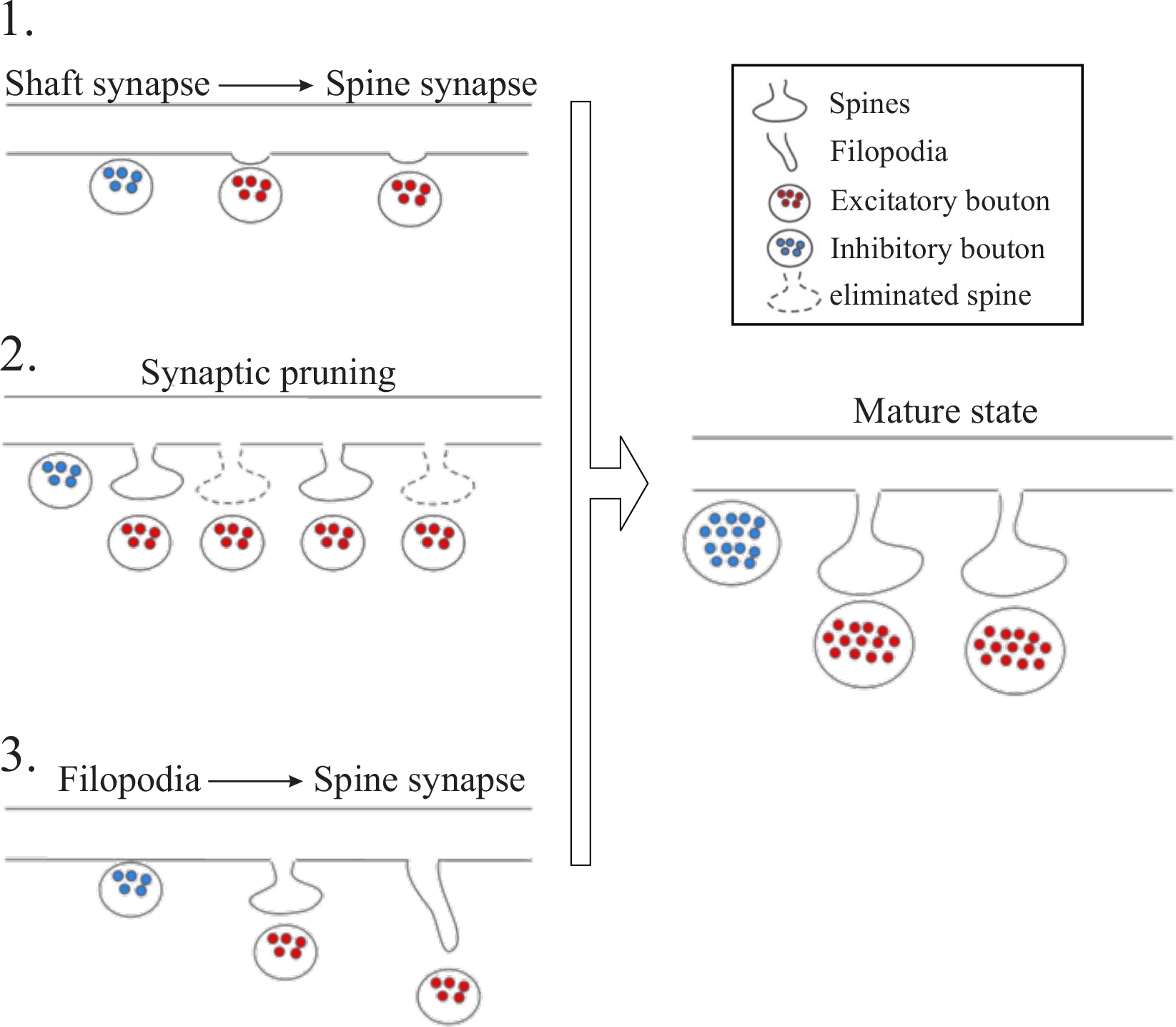
Hypothetical models of excitatory spine synapse development. *Model 1*: in neonates, some synapses on dendritic shafts bud off the dendrite to form spines (red) while others remain as shaft synapses (blue). *Model 2*: A supernumerary number of spine synapses form in neonates and subsequently are pruned. *Model 3*: neonates form filopodia extensions off dendritic shafts that mature into spine synapses.

**Supplementary Figure 13.**
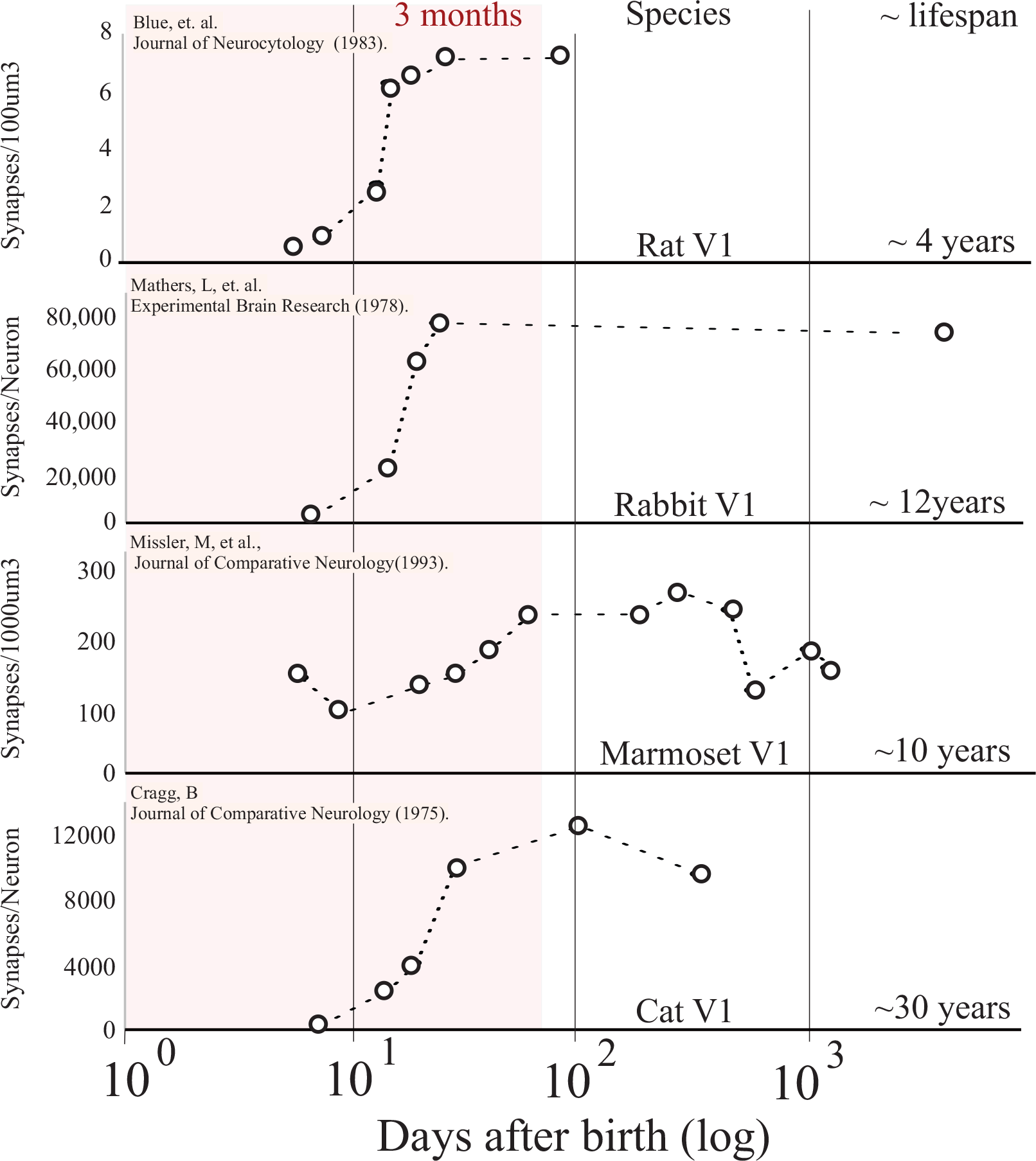
Excitatory synapse development across species. Scatter plot of x:days after birth (log) and y:published synapse density count for each listed species. For each data set, species, cortical area, average lifespan and the original publication is listed. See methods for details on how data from different publications were normalized.

**Supplementary Table 1.** Numerical averages, standard error of the mean (sem), and n number for quantification from Figure 1.

| **Figure 1b** | p7 | p75 | p3000 |
| --- | --- | --- | --- |
| Primate V1 |  |  |  |
| L2/3 | 0.31 ± 0.05,<br>n = 20, 10 µm dendrites | 2.44 ± 0.27,<br>n = 19, 10 µm dendrites | 0.86 ± 0.09,<br>n = 20, 10 µm dendrites |
| L4 | 0.65 ± 0.08,<br>n = 20, 10 µm dendrites | 2.18 ± 0.24,<br>n = 15, 10 µm dendrites | 0.32 ± 0.03,<br>n = 20, 10 µm dendrites |

|  | p6 | p14 | p36 | p105 | p523 |
| --- | --- | --- | --- | --- | --- |
| Mouse V1 |  |  |  |  |  |
| L2/3 | 0.01 ± 0.007,<br>n = 20, 10 µm dendrites | 0.87 ± 0.08,<br>n = 20, 10 µm dendrites | 1.63 ± 0.11,<br>n = 20, 10 µm dendrites | 2.06 ± 0.21,<br>n = 20, 10 µm dendrites | 1.10 ± 0.25,<br>n = 20, 10 µm dendrites |
| L4 | 0.02 ± 0.01,<br>n = 19, 10 µm dendrites | 0.77 ± 0.09,<br>n = 20, 10 µm dendrites | 0.97 ± 0.07,<br>n = 20, 10 µm dendrites | 1.74 ± 0.22,<br>n = 20, 10 µm dendrites | 1.17 ± 0.13,<br>n = 20, 10 µm dendrites |
| Mouse S1 | p9 | p14 | p60 |  |  |
| L2/3 | 0.25 ± 0.04,<br>n = 20, 10 µm dendrites | 1.21 ± 0.2,<br>n = 20, 10 µm dendrites | not done |  |  |
| L4 | 0.04 ± 0.01,<br>n = 20, 10 µm dendrites* | 0.66 ± 0.1,<br>n = 20, 10 µm dendrites | 1.87 ± 0.06,<br>n = 14, 10 µm dendrites |  |  |

\*age is p7

| Figure 1c |  | p7 |  | p75 |  | p3000 |  |  |  |  |
| --- | --- | --- | --- | --- | --- | --- | --- | --- | --- | --- |
| Primate V1 |  |  |  |  |  |  |  |  |  |  |
| L2/3 |  | 0.12 ± 0.03,<br>n = 20, 10 µm dendrites |  | 0.18 ± 0.03,<br>n = 19, 10 µm dendrites |  | 0.18 ± 0.04,<br>n = 20, 10 µm dendrites |  |  |  |  |
| L4 |  | 0.20 ± 0.03,<br>n = 20, 10 µm dendrites |  | 0.23 ± 0.03,<br>n = 15, 10 µm dendrites |  | 0.52 ± 0.08,<br>n = 20, 10 µm dendrites |  |  |  |  |
|  |  | p6 |  | p14 |  | p36 |  | p105 |  | p523 |
| Mouse V1 |  |  |  |  |  |  |  |  |  |  |
| L2/3 |  | 0.16 ± 0.03,<br>n = 20, 10 µm dendrites |  | 0.1 ± 0.02,<br>n = 20, 10 µm dendrites |  | 0.18 ± 0.03,<br>n = 20, 10 µm dendrites |  | 0.20 ± 0.03,<br>n = 20, 10 µm dendrites |  | 0.29 ± 0.04,<br>n = 20, 10 µm dendrites |
| L4 |  | 0.08 ± 0.02,<br>n = 19, 10 µm dendrites |  | 0.08 ± 0.02,<br>n = 20, 10 µm dendrites |  | 0.26 ± 0.02,<br>n = 20, 10 µm dendrites |  | 0.26 ± 0.04,<br>n = 20, 10 µm dendrites |  | 0.23 ± 0.06,<br>n = 20, 10 µm dendrites |

**Supplementary Table 2.** Numerical averages, standard error of the mean (sem), and n number for quantification from Figure 2-3.

| Figure 2-3 | excitatory axons |  |  | inhibitory axons |  |  |
| --- | --- | --- | --- | --- | --- | --- |
|  | branches/<br>um | synapses/<br>um | axons/<br>um<sup>2</sup> | synapses/<br>soma | axons/<br>soma | bouton volume (um<sup>3</sup>) |
| Primate V1,L2/3 |  |  |  |  |  |  |
| p7 | 0.16 ± 0.03,<br>n = 50 axons | 0.93 ± 0.1,<br>n = 50 axons | 5.9 ± 1.0,<br>n = 3 FOVs,<br>6 um<sup>2</sup> average area/FOV | 15.8 ± 2.1,<br>n = 95 synapses<br>across 6 soma | 13.3 ± 2.7,<br>n = 53 axons<br>across 4 soma | 0.08 ± 0.008,<br>n = 50 boutons |
| p75 | 0.38 ± 0.04,<br>n = 50 axons | 1.0 ± 0.09,<br>n = 50 axons | 4.4 ± 0.7,<br>n = 3 FOVs,<br>14 um<sup>2</sup> average area/FOV | 50.3 ± 4.1,<br>n = 302 synapses<br>across 6 soma | 22.4 ± 1.6,<br>n = 112 axons<br>across 5 soma | 0.16 ± 0.02,<br>n = 50 boutons |
| p3000 | 0.22 ± 0.03,<br>n = 50 axons | 0.46 ± 0.05,<br>n = 50 axons | 3.6 ± 0.6,<br>n = 3 FOVs,<br>13 um<sup>2</sup> average area/FOV | 16.1 ± 2.1,<br>n = 129 synapses<br>across 8 soma | 12.5 ± 1.5,<br>n = 75 axons<br>across 6 soma | 0.27 ± 0.03,<br>n = 44 boutons |
| Primate V1,L4 |  |  |  |  |  |  |
| p7 | not done | not done | not done | 23.5 ± 2.7,<br>n = 141 synapses<br>across 6 soma | not done | not done |
| p75 | not done | not done | not done | 32.7 ± 4.9,<br>n = 196 synapses<br>across 6 soma | not done | not done |
| p3000 | not done | not done | not done | 14.2 ± 1.4,<br>n = 85 synapses<br>across 6 soma | not done | not done |
|  | excitatory axons |  |  | inhibitory axons |  |  |
| Mouse V1,L2/3 | branches<br>/um | synapses/<br>um | axons/<br>um<sup>2</sup> | synapses/<br>soma * | axons/<br>soma | bouton volume (um<sup>3</sup>) |
| p6 | not done | not done | 0.83 ± 0.05,<br>n = 3 FOVs,<br>19.2 um<sup>2</sup> average area/FOV | 1.2 ± 0.3,<br>n = 7 synapses<br>across 6 soma | not done | not done |
| p14 | 0.12 ± 0.02,<br>n = 50 axons | 0.71 ± 0.09,<br>n = 50 axons | 4.1 ± 0.14,<br>n = 6 FOVs,<br>8.0 um<sup>2</sup> average area/FOV | 25.8 ± 3.5,<br>n = 155 synapses<br>across 6 soma | 17.3 ± 2.5,<br>n = 69 axons<br>across 4 soma | 0.24 ± 0.03,<br>n = 50 boutons |
| p105 | 0.22 ± 0.02,<br>n = 74 axons | 1.2 ± 0.07,<br>n = 74 axons | 4.4 ± 0.5,<br>n = 6 FOVs,<br>9.1 um<sup>2</sup> average area/FOV | 72.2 ± 5.1,<br>n = 650 synapses<br>across 9 soma | 33 ± 2.5,<br>n = 165 axons<br>across 5 soma | 0.26 ± 0.02,<br>n = 84 boutons |
| p523 | 0.15 ± 0.02,<br>n = 52 axons | 0.86 ± 0.09,<br>n = 52 axons | 4.0 ± 0.8,<br>n = 3 FOVs,<br>16 um<sup>2</sup> average area/FOV | 79.2 ± 7.1,<br>n = 475 synapses<br>across 6 soma | 39.3 ± 1.0,<br>n = 157 axons<br>across 4 soma | 0.29 ± 0.03,<br>n = 50 boutons |
| Mouse V1,L4 |  |  |  |  |  |  |
| p6 | not done | not done | not done | 2.2 ± 1.0,<br>n = 13 synapses<br>across 6 soma | not done | not done |
| p14 | not done | not done | not done | 11.3 ± 1.4,<br>n = 45 synapses<br>across 4 soma | not done | not done |
| p105 | not done | not done | not done | 42.9 ± 4.6<br>n = 45 synapses<br>across 10 soma | not done | not done |
| p523 | not done | not done | not done | 53.8 ± 7.5,<br>n = 164 synapses<br>across 6 soma | not done | not done |

\*mouse p38 synapses/soma; L2/3: $66 \pm 3.79$ , n = 396 synapses across 6 soma, L4: $38 \pm 0.82$ , n = 228 synapses across 6 soma

| Figure 4 | spine synapse density | filopodia density | synapses/soma |
| --- | --- | --- | --- |
| Mouse V1, L4 |  |  |  |
| p14 | 0.43± 0.11, n = 88 manually proofread dendrites | 0.075± 0.038, n = 88 manually proofread dendrites | 18.31± 12.15, n = 39 |
| p105 | 0.69± 0.11, n = 107 manually proofread dendrites | 0.005± 0.008, n = 107 manually proofread dendrites | 42.85± 28.8, n = 42 |

**Supplementary Table 3.** Dimensions (µm) and file size(Gb) of each dataset collected. Mouse S1 datsets can be accessed from original publication [40, 115].

| Dataset | dimensions (um) |  |  | size (Gb) |
| --- | --- | --- | --- | --- |
| Mouse V1,L2/3 |  |  |  |  |
| p6 | x: 102.8 | y: 94.5 | z: 12.0 | 81.2 |
| p14 | x: 83.7 | y: 75.3 | z: 21.1 | 92.4 |
| p38 | https://www.microns-explorer.org/phase1 |  |  |  |
| p105 | x: 107.6 | y: 99.7 | z: 30.0 | 223.66 |
| p523 | x: 111.8 | y: 95.6 | z: 22.5 | 166.82 |
| Mouse V1,L4 |  |  |  |  |
| p6 | x: 136.6 | y: 126.0 | z: 18.8 | 224.21 |
| p14 | x: 60.0 | y: 60.0 | z: 24.0 | 60.11 |
| p38 | https://www.microns-explorer.org/cortical-mm3 |  |  |  |
| p105 | x: 126.1 | y: 139.4 | z: 18.7 | 228.12 |
| p523 | x: 95.2 | y: 82.8 | z: 22.6 | 123.48 |
| Primate V1,L2/3 |  |  |  |  |
| p7 | x: 121.4 | y: 123.6 | z: 16.0 | 167.08 |
| p75 | x: 134.8 | y: 123.1 | z: 38.0 | 437.95 |
| p3000 | x: 95.3 | y: 90.9 | z: 27.0 | 162.28 |
| Primate V1,L4 |  |  |  |  |
| p7 | x: 124.4 | y: 128.7 | z: 15.5 | 172.64 |
| p75 | x: 134.8 | y: 124.9 | z: 17.9 | 209.43 |
| p3000 | x: 120.5 | y: 124.7 | z: 14.6 | 152.33 |
| Mouse S1,L2/3 |  |  |  |  |
| p9 | https://pubmed.ncbi.nlm.nih.gov/33273061/ |  |  |  |
| p14 | https://pubmed.ncbi.nlm.nih.gov/33273061/ |  |  |  |
| Mouse S1,L4 |  |  |  |  |
| p7 | https://pubmed.ncbi.nlm.nih.gov/33273061/ |  |  |  |
| p14 | https://pubmed.ncbi.nlm.nih.gov/33273061/ |  |  |  |
| p60 | https://pubmed.ncbi.nlm.nih.gov/26232230/ |  |  |  |

