## Supplementary material for "The Development of Synapses in Mouse and Macaque Primary Sensory Cortices": pairwise p-values

| Fig1b.V1 E spine syn | ms_p6 L23 | ms_p14 L23 | ms_p36_L23 | ms_p105 L23 | ms_p523 L23 | ms_p6 L4 | ms_p14 L4 | ms_p36_L4 | ms_p105 L4 | ms_p523 L4 | pr_p7 L23 | pr_p75 L23 | pr_p3000 L23 | pr_p7 L4 | pr_p75 L4 | pr_p3000 L4 |
|---|---|---|---|---|---|---|---|---|---|---|---|---|---|---|---|---|
| ms_p6 L23 | 1.00E+00 | 1.13E-08 | 1.13E-08 | 1.13E-08 | 1.33E-08 | 9.34E-02 | 1.13E-08 | 1.13E-08 | 1.13E-08 | 1.13E-08 | 2.86E-07 | 1.50E-08 | 1.13E-08 | 6.41E-08 | 5.44E-08 | 1.13E-08 |
| ms_p14 L23 | 1.13E-08 | 1.00E+00 | 4.91E-06 | 8.34E-08 | 8.41E-01 | 4.99E-08 | 3.83E-01 | 4.95E-01 | 1.14E-05 | 1.02E-01 | 1.25E-05 | 2.66E-08 | 6.40E-01 | 9.11E-02 | 1.28E-06 | 1.06E-07 |
| ms_p36_L23 | 1.13E-08 | 4.91E-06 | 1.00E+00 | 2.32E-01 | 2.58E-04 | 5.00E-08 | 9.25E-07 | 1.83E-05 | 4.61E-01 | 4.27E-03 | 9.16E-08 | 4.74E-03 | 8.18E-06 | 1.69E-07 | 3.60E-02 | 1.45E-11 |
| ms_p105 L23 | 1.13E-08 | 8.34E-08 | 2.32E-01 | 1.00E+00 | 2.49E-05 | 4.99E-08 | 3.03E-08 | 5.02E-07 | 8.10E-02 | 2.01E-04 | 6.79E-08 | 1.94E-01 | 4.07E-07 | 2.02E-09 | 3.14E-01 | 1.45E-11 |
| ms_p523 L23 | 1.33E-08 | 8.41E-01 | 2.58E-04 | 2.49E-05 | 1.00E+00 | 5.87E-08 | 6.02E-01 | 4.14E-01 | 3.30E-04 | 1.57E-01 | 3.05E-04 | 1.89E-05 | 9.25E-01 | 1.83E-01 | 1.08E-04 | 7.94E-05 |
| ms_p6 L4 | 9.34E-02 | 4.99E-08 | 5.00E-08 | 4.99E-08 | 5.87E-08 | 1.00E+00 | 4.99E-08 | 5.00E-08 | 4.99E-08 | 4.99E-08 | 3.93E-06 | 7.01E-08 | 4.99E-08 | 3.83E-07 | 3.29E-07 | 4.99E-08 |
| ms_p14 L4 | 1.13E-08 | 3.83E-01 | 9.25E-07 | 3.03E-08 | 6.02E-01 | 4.99E-08 | 1.00E+00 | 1.42E-01 | 1.65E-06 | 3.04E-02 | 5.63E-04 | 1.98E-08 | 7.99E-01 | 4.14E-01 | 3.13E-07 | 7.94E-05 |
| ms_p36_L4 | 1.13E-08 | 4.95E-01 | 1.83E-05 | 5.02E-07 | 4.14E-01 | 5.00E-08 | 1.42E-01 | 1.00E+00 | 2.14E-05 | 2.32E-01 | 2.56E-07 | 1.67E-07 | 2.01E-01 | 1.43E-02 | 6.94E-06 | 1.45E-11 |
| ms_p105 L4 | 1.13E-08 | 1.14E-05 | 4.61E-01 | 8.10E-02 | 3.30E-04 | 4.99E-08 | 1.65E-06 | 2.14E-05 | 1.00E+00 | 6.14E-03 | 7.89E-08 | 8.26E-03 | 2.40E-06 | 2.32E-08 | 2.53E-02 | 1.45E-11 |
| ms_p523 L4 | 1.13E-08 | 1.02E-01 | 4.27E-03 | 2.01E-04 | 1.57E-01 | 4.99E-08 | 3.04E-02 | 2.32E-01 | 6.14E-03 | 1.00E+00 | 7.94E-07 | 2.60E-05 | 3.04E-02 | 1.43E-03 | 3.38E-04 | 3.95E-09 |
| pr_p7 L23 | 2.86E-07 | 1.25E-05 | 9.16E-08 | 6.79E-08 | 3.05E-04 | 3.93E-06 | 5.63E-04 | 2.56E-07 | 7.89E-08 | 7.94E-07 | 1.00E+00 | 1.18E-07 | 3.98E-06 | 2.34E-03 | 1.04E-06 | 8.18E-01 |
| pr_p75 L23 | 1.50E-08 | 2.66E-08 | 4.74E-03 | 1.94E-01 | 1.89E-05 | 7.01E-08 | 1.98E-08 | 1.67E-07 | 8.26E-03 | 2.60E-05 | 1.18E-07 | 1.00E+00 | 1.31E-07 | 1.94E-09 | 6.56E-01 | 2.90E-11 |
| pr_p3000 L23 | 1.13E-08 | 6.40E-01 | 8.18E-06 | 4.07E-07 | 9.25E-01 | 4.99E-08 | 7.99E-01 | 2.01E-01 | 2.40E-06 | 3.04E-02 | 3.98E-06 | 1.31E-07 | 1.00E+00 | 2.11E-01 | 5.53E-06 | 2.32E-08 |
| pr_p7 L4 | 6.41E-08 | 9.11E-02 | 1.69E-07 | 2.02E-09 | 1.83E-01 | 3.83E-07 | 4.14E-01 | 1.43E-02 | 2.32E-08 | 1.43E-03 | 2.34E-03 | 1.94E-09 | 2.11E-01 | 1.00E+00 | 1.67E-07 | 1.77E-03 |
| pr_p75 L4 | 5.44E-08 | 1.28E-06 | 3.60E-02 | 3.14E-01 | 1.08E-04 | 3.29E-07 | 3.13E-07 | 6.94E-06 | 2.53E-02 | 3.38E-04 | 1.04E-06 | 6.56E-01 | 5.53E-06 | 1.67E-07 | 1.00E+00 | 6.16E-10 |
| pr_p3000 L4 | 1.13E-08 | 1.06E-07 | 1.45E-11 | 1.45E-11 | 7.94E-05 | 4.99E-08 | 7.94E-05 | 1.45E-11 | 1.45E-11 | 3.95E-09 | 8.18E-01 | 2.90E-11 | 2.32E-08 | 1.77E-03 | 6.16E-10 | 1.00E+00 |

| Fig1b.S1 E spine syn | ms_p9_s1_L23 | ms_p14_s1_L2 | ms_p7_s1_L4 | ms_p14_s1_L4 | ms_p105_S1_l4 |
|---|---|---|---|---|---|
| ms_p9_s1_L23 | 1.00E+00 | 1.05E-06 | 2.81E-05 | 1.78E-03 | 1.05E-06 |
| ms_p14_s1_L23 | 1.05E-06 | 1.00E+00 | 4.49E-08 | 3.89E-03 | 1.36E-03 |
| ms_p7_s1_L4 | 2.81E-05 | 4.49E-08 | 1.00E+00 | 1.51E-07 | 5.96E-07 |
| ms_p14_s1_L4 | 2.81E-05 | 4.49E-08 | 1.00E+00 | 1.00E+00 | 2.11E-06 |

| Fig1c. V1 E shaft syn | ms_p6 L23 | ms_p14 L23 | ms_p36_L23 | ms_p105 L23 | ms_p523 L23 | ms_p6 L4 | ms_p14 L4 | ms_p36_L4 | ms_p105 L4 | ms_p523 L4 | pr_p7 L23 | pr_p75 L23 | pr_p3000 L23 | pr_p7 L4 | pr_p75 L4 | pr_p3000 L4 |
|---|---|---|---|---|---|---|---|---|---|---|---|---|---|---|---|---|
| ms_p6 L23 | 1.00E+00 | 8.91E-02 | 5.07E-01 | 6.14E-01 | 4.99E-02 | 1.60E-02 | 1.60E-02 | 3.06E-03 | 5.63E-02 | 6.16E-01 | 1.13E-01 | 5.27E-01 | 9.46E-01 | 3.23E-01 | 3.72E-02 | 1.44E-04 |
| ms_p14 L23 | 8.91E-02 | 1.00E+00 | 1.61E-02 | 3.60E-01 | 5.44E-03 | 5.49E-01 | 3.79E-01 | 9.24E-06 | 1.07E-03 | 5.75E-02 | 9.02E-01 | 6.01E-02 | 1.07E-01 | 7.37E-03 | 4.30E-04 | 2.89E-06 |
| ms_p36_L23 | 5.07E-01 | 1.61E-02 | 1.00E+00 | 3.50E-01 | 1.20E-01 | 5.05E-03 | 5.67E-03 | 9.78E-03 | 1.48E-01 | 9.89E-01 | 3.57E-02 | 8.77E-01 | 4.23E-01 | 9.03E-01 | 1.13E-01 | 3.05E-04 |
| ms_p105 L23 | 6.14E-01 | 3.60E-01 | 3.50E-01 | 1.00E+00 | 3.42E-02 | 1.85E-01 | 5.85E-02 | 1.55E-02 | 7.72E-02 | 4.58E-01 | 4.66E-01 | 4.23E-01 | 8.95E-01 | 2.06E-01 | 9.67E-02 | 8.68E-05 |
| ms_p523 L23 | 4.99E-02 | 5.44E-03 | 1.20E-01 | 3.42E-02 | 1.00E+00 | 2.70E-04 | 4.01E-04 | 1.00E+00 | 6.55E-01 | 2.39E-01 | 2.33E-03 | 1.73E-01 | 9.57E-02 | 4.25E-01 | 6.09E-01 | 2.63E-02 |
| ms_p6 L4 | 1.60E-02 | 5.49E-01 | 5.05E-03 | 1.85E-01 | 2.70E-04 | 1.00E+00 | 5.03E-01 | 2.15E-06 | 1.06E-03 | 2.46E-02 | 3.89E-01 | 6.46E-03 | 8.05E-02 | 2.03E-03 | 2.79E-04 | 1.41E-06 |
| ms_p14 L4 | 1.60E-02 | 3.79E-01 | 5.67E-03 | 5.85E-02 | 4.01E-04 | 5.03E-01 | 1.00E+00 | 3.07E-05 | 6.73E-04 | 1.50E-02 | 2.49E-01 | 1.04E-02 | 7.85E-02 | 4.39E-03 | 8.56E-04 | 6.91E-06 |
| ms_p36_L4 | 3.06E-03 | 9.24E-06 | 9.78E-03 | 1.55E-02 | 1.00E+00 | 2.15E-06 | 3.07E-05 | 1.00E+00 | 9.68E-01 | 9.60E-02 | 2.90E-05 | 1.63E-02 | 8.93E-03 | 3.60E-02 | 4.99E-01 | 1.55E-02 |
| ms_p105 L4 | 5.63E-02 | 1.07E-03 | 1.48E-01 | 7.72E-02 | 6.55E-01 | 1.06E-03 | 6.73E-04 | 9.68E-01 | 1.00E+00 | 3.09E-01 | 4.22E-03 | 1.48E-01 | 5.18E-02 | 1.98E-01 | 6.65E-01 | 2.94E-02 |
| ms_p523 L4 | 6.16E-01 | 5.75E-02 | 9.89E-01 | 4.58E-01 | 2.39E-01 | 2.46E-02 | 1.50E-02 | 9.60E-02 | 3.09E-01 | 1.00E+00 | 8.96E-02 | 8.77E-01 | 4.37E-01 | 7.35E-01 | 2.64E-01 | 2.79E-03 |
| pr_p7 L23 | 1.13E-01 | 9.02E-01 | 3.57E-02 | 4.66E-01 | 2.33E-03 | 3.89E-01 | 2.49E-01 | 2.90E-05 | 4.22E-03 | 8.96E-02 | 1.00E+00 | 3.13E-02 | 1.52E-01 | 7.61E-03 | 2.52E-03 | 1.24E-05 |
| pr_p75 L23 | 5.27E-01 | 6.01E-02 | 8.77E-01 | 4.23E-01 | 1.73E-01 | 6.46E-03 | 1.04E-02 | 1.63E-02 | 1.48E-01 | 8.77E-01 | 3.13E-02 | 1.00E+00 | 7.24E-01 | 5.46E-01 | 1.36E-01 | 5.77E-04 |
| pr_p3000 L23 | 9.46E-01 | 1.07E-01 | 4.23E-01 | 8.95E-01 | 9.57E-02 | 8.05E-02 | 7.85E-02 | 8.93E-03 | 5.18E-02 | 4.37E-01 | 1.52E-01 | 7.24E-01 | 1.00E+00 | 5.23E-01 | 3.65E-02 | 2.68E-04 |
| pr_p7 L4 | 3.23E-01 | 7.37E-03 | 9.03E-01 | 2.06E-01 | 4.25E-01 | 2.03E-03 | 4.39E-03 | 3.60E-02 | 1.98E-01 | 7.35E-01 | 7.61E-03 | 5.46E-01 | 5.23E-01 | 1.00E+00 | 1.99E-01 | 7.57E-04 |
| pr_p75 L4 | 3.72E-02 | 4.30E-04 | 1.13E-01 | 9.67E-02 | 6.09E-01 | 2.79E-04 | 8.56E-04 | 4.99E-01 | 6.65E-01 | 2.64E-01 | 2.52E-03 | 1.36E-01 | 3.65E-02 | 1.99E-01 | 1.00E+00 | 1.43E-02 |
| pr_p3000 L4 | 1.44E-04 | 2.89E-06 | 3.05E-04 | 8.68E-05 | 2.63E-02 | 1.41E-06 | 6.91E-06 | 1.55E-02 | 2.94E-02 | 2.79E-03 | 1.24E-05 | 5.77E-04 | 2.68E-04 | 7.57E-04 | 1.43E-02 | 1.00E+00 |

| Fig2b. V1 E axon syn per um | ms_p14_l23 | ms_p105_l23 | ms_p523_l23 | pr_p7_l23 | pr_p75_l23 | pr_p3000_l23 |
|---|---|---|---|---|---|---|
| ms_p14_l23 | 1.00E+00 | 8.49E-09 | 2.93E-02 | 7.93E-02 | 1.77E-03 | 1.11E-02 |
| ms_p105_l23 | 8.49E-09 | 1.00E+00 | 1.61E-05 | 0.008139, | 3.65E-02 | 1.44E-14 |
| ms_p523_l23 | 2.93E-02 | 1.61E-05 | 1.00E+00 | 6.76E-01 | 1.21E-01 | 3.23E-06 |
| pr_p7_l23 | 7.93E-02 | 0.008139, | 6.76E-01 | 1.00E+00 | 3.76E-01 | 7.39E-04 |
| pr_p75_l23 | 1.77E-03 | 3.65E-02 | 1.21E-01 | 3.76E-01 | 1.00E+00 | 2.66E-07 |
| pr_p3000_l23 | 1.11E-02 | 1.44E-14 | 3.23E-06 | 7.39E-04 | 2.66E-07 | 1.00E+00 |

| Fig 2c. V1 E axon branches per | ms_p14_l23 | ms_p105_l23 | ms_p523_l23 | pr_p7_l23 | pr_p75_l23 | pr_p3000_l23 |
|---|---|---|---|---|---|---|
| ms_p14_l23 | 1.00E+00 | 7.29E-03 | 5.38E-01 | 9.46E-01 | 1.05E-06 | 2.01E-02 |
| ms_p105_l23 | 7.29E-03 | 1.00E+00 | 6.97E-02 | 2.11E-02 | 2.11E-03 | 9.96E-01 |
| ms_p523_l23 | 5.38E-01 | 6.97E-02 | 1.00E+00 | 5.98E-01 | 1.42E-05 | 1.15E-01 |
| pr_p7_l23 | 9.46E-01 | 2.11E-02 | 5.98E-01 | 1.00E+00 | 1.93E-05 | 5.90E-02 |
| pr_p75_l23 | 1.05E-06 | 2.11E-03 | 1.42E-05 | 1.93E-05 | 1.00E+00 | 4.74E-03 |
| pr_p3000_l23 | 2.01E-02 | 9.96E-01 | 1.15E-01 | 5.90E-02 | 4.74E-03 | 1.00E+00 |

| Fig 3b. V1 syn per soma | ms_p6 L23 | ms_p14 L23 | ms_p36_L23 | ms_p105 L23 | ms_p523 L23 | ms_p6 L4 | ms_p14 L4 | ms_p36_L4 | ms_p105 L4 | ms_p523 L4 | pr_p7 L23 | pr_p75 L23 | pr_p3000 L23 | pr_p7 L4 | pr_p75 L4 | pr_p3000 L4 |
|---|---|---|---|---|---|---|---|---|---|---|---|---|---|---|---|---|
| ms_p6 L23 | 1.00E+00 | 4.70E-03 | 4.70E-03 | 1.69E-03 | 4.70E-03 | 7.97E-01 | 1.28E-02 | 4.62E-03 | 9.52E-03 | 4.70E-03 | 4.70E-03 | 4.70E-03 | 2.23E-03 | 4.62E-03 | 4.55E-03 | 4.55E-03 |
| ms_p14 L23 | 4.70E-03 | 1.00E+00 | 2.17E-03 | 1.77E-03 | 2.16E-03 | 4.77E-03 | 9.52E-03 | 1.01E-02 | 4.80E-02 | 8.73E-01 | 4.46E-02 | 2.16E-03 | 2.31E-02 | 5.75E-01 | 3.77E-01 | 7.69E-03 |
| ms_p36_L23 | 4.70E-03 | 2.17E-03 | 1.00E+00 | 4.09E-01 | 1.99E-01 | 4.77E-03 | 9.52E-03 | 5.00E-03 | 2.86E-03 | 3.36E-01 | 2.17E-03 | 6.49E-02 | 2.36E-03 | 5.00E-03 | 4.92E-03 | 4.92E-03 |
| ms_p105 L23 | 1.69E-03 | 1.77E-03 | 4.09E-01 | 1.00E+00 | 5.95E-01 | 1.71E-03 | 6.85E-03 | 1.76E-03 | 2.89E-04 | 3.16E-03 | 1.77E-03 | 2.11E-02 | 6.21E-04 | 1.76E-03 | 1.74E-03 | 1.74E-03 |
| ms_p523 L23 | 4.70E-03 | 2.16E-03 | 1.99E-01 | 5.95E-01 | 1.00E+00 | 4.77E-03 | 9.52E-03 | 5.00E-03 | 1.51E-03 | 1.03E-02 | 2.16E-03 | 2.60E-02 | 2.36E-03 | 5.00E-03 | 4.92E-03 | 4.92E-03 |
| ms_p6 L4 | 7.97E-01 | 4.77E-03 | 4.77E-03 | 1.71E-03 | 4.77E-03 | 1.00E+00 | 1.31E-02 | 4.70E-03 | 9.96E-05 | 7.80E-03 | 4.77E-03 | 4.77E-03 | 2.26E-03 | 4.70E-03 | 4.62E-03 | 4.62E-03 |
| ms_p14 L4 | 1.28E-02 | 9.52E-03 | 9.52E-03 | 6.85E-03 | 9.52E-03 | 1.31E-02 | 1.00E+00 | 1.39E-02 | 3.58E-03 | 1.14E-01 | 1.71E-01 | 9.52E-03 | 1.04E-01 | 1.39E-02 | 1.36E-02 | 1.59E-01 |
| ms_p36_L4 | 4.62E-03 | 1.01E-02 | 5.00E-03 | 1.76E-03 | 5.00E-03 | 4.70E-03 | 1.39E-02 | 1.00E+00 | 3.98E-01 | 1.71E-01 | 5.00E-03 | 6.30E-03 | 2.34E-03 | 4.92E-03 | 3.76E-01 | 4.85E-03 |
| ms_p105 L4 | 9.32E-05 | 4.80E-02 | 2.86E-03 | 2.89E-04 | 1.51E-03 | 9.96E-05 | 3.58E-03 | 3.98E-01 | 1.00E+00 | 9.70E-02 | 3.20E-03 | 2.56E-02 | 1.98E-04 | 5.25E-03 | 2.32E-01 | 6.80E-04 |
| ms_p523 L4 | 4.70E-03 | 8.73E-01 | 3.36E-01 | 3.16E-03 | 1.03E-02 | 7.80E-03 | 1.14E-01 | 1.71E-01 | 9.70E-02 | 1.00E+00 | 3.10E-01 | 3.70E-02 | 2.44E-01 | 1.00E+00 | 2.28E-01 | 1.47E-01 |
| pr_p7 L23 | 4.70E-03 | 4.46E-02 | 2.17E-03 | 1.77E-03 | 2.16E-03 | 4.77E-03 | 1.71E-01 | 5.00E-03 | 3.20E-03 | 3.10E-01 | 1.00E+00 | 2.16E-03 | 1.00E+00 | 1.08E-01 | 8.02E-03 | 6.27E-01 |
| pr_p75 L23 | 4.70E-03 | 2.16E-03 | 6.49E-02 | 2.11E-02 | 2.60E-02 | 4.77E-03 | 9.52E-03 | 6.30E-03 | 2.56E-02 | 3.70E-02 | 2.16E-03 | 1.00E+00 | 2.36E-03 | 5.00E-03 | 6.46E-02 | 4.92E-03 |
| pr_p3000 L23 | 2.23E-03 | 2.31E-02 | 2.36E-03 | 6.21E-04 | 2.36E-03 | 2.26E-03 | 1.04E-01 | 2.34E-03 | 1.98E-04 | 2.44E-01 | 1.00E+00 | 2.36E-03 | 1.00E+00 | 9.08E-02 | 1.15E-02 | 6.45E-01 |
| pr_p7 L4 | 4.62E-03 | 5.75E-01 | 5.00E-03 | 1.76E-03 | 5.00E-03 | 4.70E-03 | 1.39E-02 | 4.92E-03 | 5.25E-03 | 1.00E+00 | 1.08E-01 | 5.00E-03 | 9.08E-02 | 1.00E+00 | 4.20E-01 | 6.32E-02 |
| pr_p75 L4 | 4.55E-03 | 3.77E-01 | 4.92E-03 | 1.74E-03 | 4.92E-03 | 4.62E-03 | 1.36E-02 | 3.76E-01 | 2.32E-01 | 2.28E-01 | 8.02E-03 | 6.46E-02 | 1.15E-02 | 4.20E-01 | 1.00E+00 | 4.77E-03 |
| pr_p3000 L4 | 4.55E-03 | 7.69E-03 | 4.92E-03 | 1.74E-03 | 4.92E-03 | 4.62E-03 | 1.59E-01 | 4.85E-03 | 6.80E-04 | 1.47E-01 | 6.27E-01 | 4.92E-03 | 6.45E-01 | 6.32E-02 | 4.77E-03 | 1.00E+00 |

| Fig 3c. V1 axons per soma | ms_p14_l23 | ms_p105_l23 | ms_p523_l23 | pr_p7_l23 | pr_p75_l23 | pr_p3000_l23 |
|---|---|---|---|---|---|---|
| ms_p14_l23 | 1.00E+00 | 1.59E-02 | 2.94E-02 | 1.89E-01 | 1.38E-01 | 1.98E-01 |
| ms_p105_l23 | 1.59E-02 | 1.00E+00 | 1.10E-01 | 1.59E-02 | 2.12E-02 | 7.97E-03 |
| ms_p523_l23 | 2.94E-02 | 1.10E-01 | 1.00E+00 | 2.94E-02 | 1.89E-02 | 1.36E-02 |
| pr_p7_l23 | 1.89E-01 | 1.59E-02 | 2.94E-02 | 1.00E+00 | 4.81E-02 | 9.14E-01 |
| pr_p75_l23 | 1.38E-01 | 2.12E-02 | 1.89E-02 | 4.81E-02 | 1.00E+00 | 1.33E-02 |
| pr_p3000_l23 | 1.98E-01 | 7.97E-03 | 1.36E-02 | 9.14E-01 | 1.33E-02 | 1.00E+00 |

| Fig 3e. V1 soma syn bouton volume | ms_p14_l23 | ms_p105_l23 | ms_p523_l23 | pr_p7_l23 | pr_p75_l23 | pr_p3000_l23 |
|---|---|---|---|---|---|---|
| ms_p14_l23 | 1.00E+00 | 7.56E-01 | 1.45E-01 | 1.40E-08 | 6.71E-03 | 4.64E-01 |
| ms_p105_l23 | 7.56E-01 | 1.00E+00 | 2.39E-01 | 2.58E-12 | 6.77E-05 | 6.94E-01 |
| ms_p523_l23 | 1.45E-01 | 2.39E-01 | 1.00E+00 | 3.65E-11 | 2.76E-05 | 5.06E-01 |
| pr_p7_l23 | 1.40E-08 | 2.58E-12 | 3.65E-11 | 1.00E+00 | 3.70E-05 | 3.76E-09 |
| pr_p75_l23 | 6.71E-03 | 6.77E-05 | 2.76E-05 | 3.70E-05 | 1.00E+00 | 5.11E-04 |
| pr_p3000_l23 | 4.64E-01 | 6.94E-01 | 5.06E-01 | 3.76E-09 | 5.11E-04 | 1.00E+00 |

| Key | description | stats test |
| --- | --- | --- |
| Fig1b.V1 E spine syn | excitatory spine synpase/µm quantification across 10 µm dendrite fragments in mouse and primate [illegible] | Mann Whitney U |
| Fig1b.S1 E spine syn | excitatory spine synpase/µm quantification across 10 µm dendrite fragments in mouse S1 | Mann Whitney U |
| Fig1c. V1 E shaft syn | inhibitory shaft synpase/µm quantification across 10 µm dendrite fragments in mouse and primate V | Mann Whitney U |
| Fig2b. V1 E axon syn per um | synapses/µm of excitatory axons in mouse and primate V1 | Mann Whitney U |
| Fig 2c. V1 E axon branches per um | branches/µm of excitatory axons in mouse and primate V1 | Mann Whitney U |
| Fig 3b. V1 syn per soma | total number of synapses on excitatory some in mouse and primate V1 | Mann Whitney U |
| Fig 3c. V1 axons per soma | total number of axons that synapses onto excitatory some in mouse and primate V1 | Mann Whitney U |
| Fig 3e. V1 soma syn bouton volume | bouton volume of soma synapses in mouse and primate V1 | Mann Whitney U |
