## supplemental species data for "The Development of Synapses in Mouse and Macaque Primary Sensory Cortices"

https://onlinelibrary.wiley.com/doi/epdf/10.1002/cne.901600202

| | | |
|---|---|---|
| species | cat Visual cortex | |
| ave lifespan | 12.5 years | |
| postnatal da | estimate val | %, p7.5 set to 0 |
| 7.5 | 250 | 0 |
| 15 | 2500 | 10 |
| 20 | 4000 | 16 |
| 30 | 10000 | 40 |
| 108 | 12500 | 50 |
| 375 | 9500 | 38 |

https://link.springer.com/content/pdf/10.1007/BF01181531.pdf

| | | |
|---|---|---|
| species | Rat Visual cortex | |
| ave lifespan | black = 12 months, brown = 48 months | |
| postnatal da | estimate val | %, p6 set to 0 |
| 6 | 0.677 | 0 |
| 8 | 0.894 | 1.32053176 |
| 14 | 2.7 | 3.98818316 |
| 16 | 6.235 | 9.20974889 |
| 20 | 6.4 | 9.4534712 |
| 28 | 7.132 | 10.534712 |
| 90 | 7.169 | 10.5893648 |

https://onlinelibrary.wiley.com/doi/epdf/10.1002/cne.903330105

| | | |
|---|---|---|
| species | marmoset v1 | |
| ave lifespan | black = 12 years | |
| postnatal da | estimate val | %, p6 set to 0 |
| 6 | 127 | 0 |
| 9 | 81 | 0.63779528 |
| 20 | 118 | 0.92913386 |
| 28 | 131 | 1.03149606 |
| 40 | 164 | 1.29133858 |
| 60 | 215 | 1.69291339 |
| 180 | 214 | 1.68503937 |
| 270 | 245 | 1.92913386 |
| 450 | 220 | 1.73228346 |
| 570 | 111 | 0.87401575 |
| 1020 | 166 | 1.30708661 |
| 1260 | 141 | 1.11023622 |

https://pubmed.ncbi.nlm.nih.gov/729659/

| | | | |
|---|---|---|---|
| | species | rabbit v1 | |
| | ave lifespan | 9 years | |
| | postnatal da | estimate val | %, p6 set to 0 |
| | 7 | 250 | 0 |
| | 15 | 2200 | 8.8 |
| | 20 | 6300 | 25.2 |
| | 25 | 7800 | 31.2 |
| adult | 3240 | 7400 | 29.6 |
